# A novel class of TMPRSS2 inhibitors potently block SARS-CoV-2 and MERS-CoV viral entry and protect human epithelial lung cells

**DOI:** 10.1101/2021.05.06.442935

**Authors:** Matthew Mahoney, Vishnu C. Damalanka, Michael A. Tartell, Dong Hee Chung, André Luiz Lourenco, Dustin Pwee, Anne E. Mayer Bridwell, Markus Hoffmann, Jorine Voss, Partha Karmakar, Nurit Azouz, Andrea M. Klingler, Paul W. Rothlauf, Cassandra E. Thompson, Melody Lee, Lidija Klampfer, Christina Stallings, Marc E. Rothenberg, Stefan Pöhlmann, Sean P. Whelan, Anthony J. O’Donoghue, Charles S. Craik, James W. Janetka

**Affiliations:** Department of Biochemistry and Molecular Biophysics, Washington University School of Medicine, Saint Louis, Missouri, United States; ProteXase Therapeutics, Inc., Saint Louis, Missouri, United States; Department of Molecular Microbiology, Washington University School of Medicine, Saint Louis, Missouri, United States; Program in Virology, Harvard Medical School, Boston, Massachusetts, United States; Department of Pharmaceutical Chemistry, University of California, San Francisco, California, United States; Skaggs School of Pharmacy and Pharmaceutical Sciences, University of California, San Diego, California, United States; Infection Biology Unit, German Primate Center, Leibniz Institute for Primate Research, Göttingen, Germany; Division of Allergy and Immunology, Cincinnati Children’s Hospital Medical Center and Department of Pediatrics, University of Cincinnati College of Medicine, Cincinnati, Ohio, United States; Faculty of Biology and Psychology, Georg-August University Göttingen, Göttingen, Germany

## Abstract

The host cell serine protease TMPRSS2 is an attractive therapeutic target for COVID-19 drug discovery. This protease activates the Spike protein of Severe Acute Respiratory Syndrome Coronavirus 2 (SARS-CoV-2) and of other coronaviruses and is essential for viral spread in the lung. Utilizing rational structure-based drug design (SBDD) coupled to substrate specificity screening of TMPRSS2, we have discovered a novel class of small molecule ketobenzothiazole TMPRSS2 inhibitors with significantly improved activity over existing irreversible inhibitors Camostat and Nafamostat. Lead compound MM3122 (**4**) has an IC_50_ of 340 pM against recombinant full-length TMPRSS2 protein, an EC_50_ of 430 pM in blocking host cell entry into Calu-3 human lung epithelial cells of a newly developed VSV SARS-CoV-2 chimeric virus, and an EC_50_ of 74 nM in inhibiting cytopathic effects induced by SARS-CoV-2 virus in Calu-3 cells. Further, MM3122 blocks Middle East Respiratory Syndrome Coronavirus (MERS-CoV) cell entry with an EC_50_ of 870 pM. MM3122 has excellent metabolic stability, safety, and pharmacokinetics in mice with a half-life of 8.6 hours in plasma and 7.5 h in lung tissue, making it suitable for in vivo efficacy evaluation and a promising drug candidate for COVID-19 treatment.

## Introduction

Severe Acute Respiratory Syndrome Coronavirus-2 (SARS-CoV-2) is the newly emerged, highly transmissible coronavirus responsible for the ongoing Coronavirus Disease 2019 (COVID-19) pandemic, which is associated with 136 million cases and almost 3 million deaths worldwide as of April 12, 2021 (https://coronavirus.jhu.edu/map.html). While three vaccines have recently been approved by the FDA, there are still no clinically approved small molecule drugs available for the treatment of this disease except Remdesivir and the effectiveness of the vaccines against immune escape variants might be reduced. Multiple therapeutic strategies have been proposed^*2, 3*^, including both viral and host proteins but none have yet been fully validated for clinical application. One class of protein targets which have shown promising results are proteolytic enzymes including the viral proteases^*2, 4–6*^, Papain-Like Protease (PLpro) and the 3C-like or ‘Main Protease’ (3CL or MPro), and several host proteases involved in viral entry, replication, and effects on the immune system creating the life-threatening symptoms of COVID-19 infection.^*5–7*^. The latter include various members of the cathepsin family of cysteine proteases including cathepsin L, furin, and the serine proteases factor Xa, plasmin, elastase, tryptase, TMPRSS2 and TMPRSS4.

TMPRSS2^*8*^ is a type II transmembrane serine proteases (TTSP)^*9*^ that has been shown to be crucial for host-cell viral entry and spread of SARS-CoV-2^*10–12*^, as well as SARS-CoV^*13, 14*^, Middle East Respiratory Syndrome Coronavirus (MERS-CoV)^*15*^ and influenza A viruses^*16–23*^. Like SARS-CoV and MERS-CoV, SARS-CoV-2 cell entry involves binding of the viral Spike protein to the host cell receptor Angiotensin Converting Enzyme-2 (ACE2). The Spike protein requires proteolytic processing/priming by TMPRSS2 to mediate entry into lung cells, thus small molecule inhibitors of this target offer much promise as new therapeutics for COVID-19 and other coronavirus diseases^*10, 11*^. It has been demonstrated that the TMPRSS2-expressing lung epithelial Calu-3 cells are highly permissive to SARS-CoV-2 infection. The irreversible serine protease inhibitors Camostat^*10*^ and Nafamostat^*24*^ are effective at preventing host cell entry and replication of SARS-CoV-2 in Calu-3 cells through a TMPRSS2-dependent mechanism.

Herein, we report on the discovery of a new class of substrate-based ketobenzothiazole (kbt) inhibitors of TMPRSS2 with potent antiviral activity against SARS-CoV-2 and which are significantly improved over Camostat and Nafamostat. Several compounds were found to be strong inhibitors of viral entry and replication, with EC_50_ values exceeding the potency of Camostat and Nafamostat and without cytotoxicity. Newly developed compound MM3122 (**4**) has excellent pharmacokinetics (PK) and safety in mice and is thus a promising new lead candidate drug for COVID-19 treatment.

## Results and Discussion

### 1. Hit identification of TMPRSS2 inhibitors

It is well established that several TTSPs including TMPRSS2 not only play a role in infectious diseases^*8*^, but also cancer progression and metastasis^*25–27*^ which is thought to be mainly through its ability to activate hepatocyte growth factor (HGF), the sole ligand for MET receptor tyrosine kinase. This is accomplished via proteolytic processing of the inactive single-chain precursor pro-HGF to a two-chain active form. TMPRSS2 shares pro-HGF as a protein substrate with the other HGF-activating serine proteases, HGFA (HGF-Activator), hepsin and matriptase.^*28, 29*^ Like TMPRSS2, other TTSPs such as matriptase and hepsin have a canonical serine protease domain residing as the C-terminal domain of the protein that is anchored to the cell membrane by an N-terminal type II signal peptide thereby presenting their enzymatic activity outside the cell. We have previously reported on the discovery and anticancer properties of peptidomimetic ketothiazole (kt) and ketobenzothiazole (kbt) inhibitors of HGFA, matriptase and hepsin^*30–34*^ named synthetic HGF Activation Inhibitors or sHAIs. Since TMPRSS2 has an overlapping endogenous substrate specificity profile with HGF-activating proteases, we postulated that our substrate-based sHAIs would also inhibit TMPRSS2.

#### Antiviral activity of sHAIs (VSV/SARS-CoV-2 Spike Protein Pseudotypes)

Based on our premise that sHAIs would inhibit TMPRSS2, we first selected the two sHAI lead compounds, ZFH7116 (**1**)^*32, 35*^ and VD2173 (**2**)^*36*^ (**Figure 1**) for initial antiviral testing and confirmed that they potently inhibit TMPRSS2-dependent host cell viral entry driven by the spike protein of SARS-CoV-2 (**SARS-2 S, Figure 2A**) into Calu-3 lung epithelial cells having an EC_50_ of 307 and 104 nM, respectively. Marked inhibition of SARS-2-S-driven entry was also detected for the irreversible inhibitor Camostat, in line with published data^*10, 24*^. It is noteworthy that neither compound showed activity in TMPRSS2 negative Vero cells or modulated entry into Calu-3 cells by pseudotypes bearing the VSV glycoprotein (VSVpp VSV-G) (Supplementary data) suggesting that the inhibition of SARS-CoV-2 spike-driven entry into Calu-3 cells was due to blockade of TMPRSS2.

**Figure 1.**
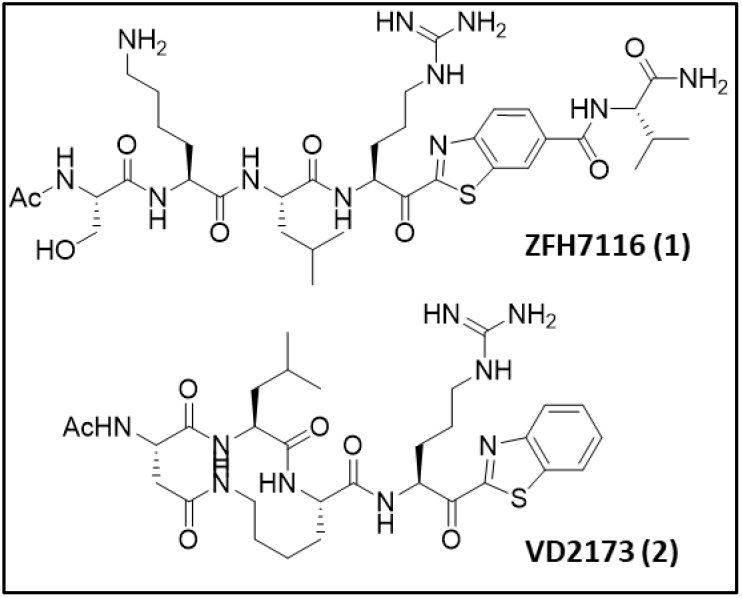
Structures of the initial sHAI ketobenzothiazole (kbt) inhibitors of TMPRSS2.

**Figure 2.**
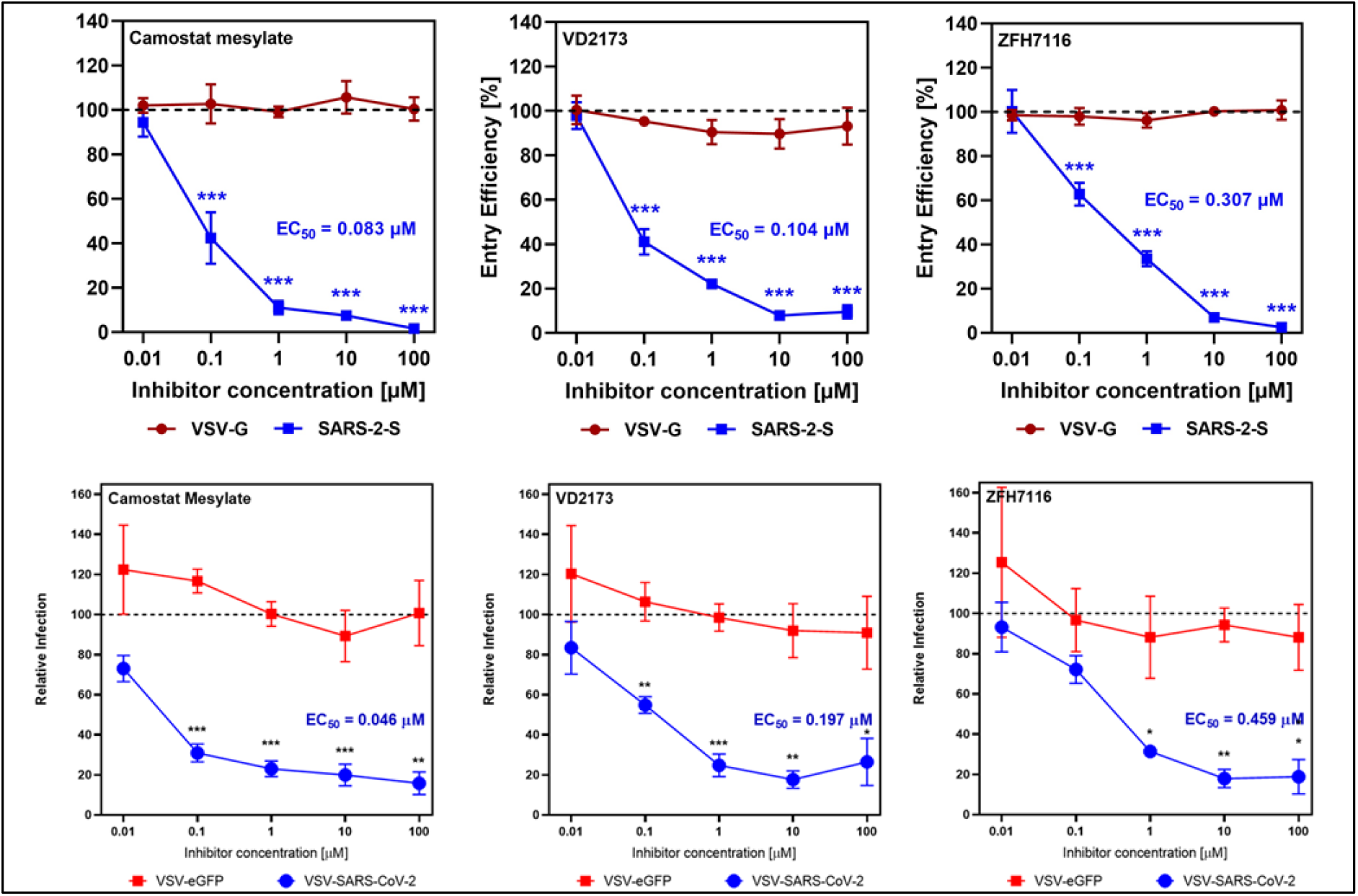
Inhibition of SARS-CoV-2 cell entry into Calu-3 lung epithelial cells by ZFH7116 (1) and VD2173 (2) using. A) VSV-SARS-CoV2-Spike protein Pseudotypes and B) VSV-SARS-CoV2-Spike protein Chimeras. EC_50_s are calculated from an average of 3 separate experiments. Camostat, was used as a positive control.

#### Antiviral activity of sHAIs (VSV/SARS-CoV-2 Spike Protein Chimeras)

Using a replication-competent, chimeric VSV expressing the SARS-CoV-2 Spike (shown to mimic the Spike-dependent entry of authentic SARS-CoV-2^*37*^ we also demonstrated that **1** and **2** block viral entry into Calu-3 cells in a dose-dependent manner. Both drugs displayed EC_50_ values of 459 and 197 nM) against the chimeric virus VSV-SARS-CoV-2 (**Figure 2B**) compared to the pseudotyped virus (**Figure 2A**), while showing no activity against VSV-eGFP (VSV G dependent entry), or against either virus in Vero cells (Supplementary data), which are TMPRSS2 negative. This confirms our initial result and allows for the establishment of a system for screening antiviral activity of these inhibitors using VSV-SARS-CoV-2.

#### Inhibition of cell bound TMPRS2 enzymatic activity by sHAIs

We tested the inhibitory activity of **1** and **2** on TMPRSS2 proteolytic activity in a cell-based enzyme-based fluorogenic assay, by overexpression of TMPRSS2 in a human cell line, HEK-293T, commonly used for experimentation because of its high transfectability. As previously demonstrated, expression of TMPRSS2 can be accurately measured^*38, 39*^ using a fluorogenic peptide reporter substrate, Boc-QAR-AMC in cell cultures. **1** inhibited cell based TMPRSS2 enzyme activity in a concentration dependent manner between 10 μM to 10 nM with an IC_50_ of 314 nM (Supplementary data). **2** was a more potent inhibitor of TMPRSS2 proteolytic activity with IC_50_ of 57 nM. These data confirm that both **1** and **2** mediate their function by potently inhibiting TMPRSS2.

#### Selectivity data of TMPRSS2 inhibitors in a panel of proteases

We profiled **1** and **2** for their selectivity against a panel of 43 serine and cysteine proteases (Supplementary material) for comparison to that of Camostat and Nafamostat protease selectivity data which has been published^*40*^. We found **1** was a potent inhibitor of matriptase-2, plasma kallikrein, proteinase K, trypsin, tryptase b2 and G1 but also was a moderate inhibitor of factor Xa, factor XIIa, kallikrein 5 and 14 as well as cysteine protease cathepsin S while showing some activity against cathepsin L as well. There was overlap of **1** with the selectivity profiles of Camostat except they did not show any activity against the cathepsins. Furthermore, Camostat more potently inhibited factor XIIa, plasma kallikrein, matriptase-2, and plasmin while also having increased activity for trypsin, the tryptases and urokinase. Cyclic peptide **2** also more potently inhibited urokinase and factor XIIa relative to **1** but not cathepsin S or L, and instead was a moderate inhibitor of cathepsin B.

The ramifications of these different selectivity profiles may explain the varied activity in the VSV-pseudotype and chimeric viral entry results most notably the urokinase potency for which **1** was significantly weaker.

### 2. Lead identification of TMPRSS2 inhibitors

We published^*33*^ that Camostat and Nafamostat (**Figure 3B**) are inhibitors of matriptase and hepsin, and analog Nafamostat is the most active published inhibitor of SARS-CoV-2 cell entry^*11*^. While we previously pursued this chemical series as HGFA, matriptase and hepsin inhibitors^*33*^, the best series we have developed are small peptide-based molecules like **1, 2** and **3** (Ac-SKLR-kbt; **Figure 3B**) which exhibit low nM to picomolar IC_50_s. These compounds contain a serine trapping ketobenzothiazole (kbt) warhead^*30–32, 34*^ which reacts covalently with the protease, but importantly in a reversible manner, unlike Camostat or Nafamostat in which the inhibition is irreversible. Therefore, we pursued the kbt class of inhibitors for lead identification studies towards more potent and selective TMPRSS2 inhibitors.

**Figure 3.**
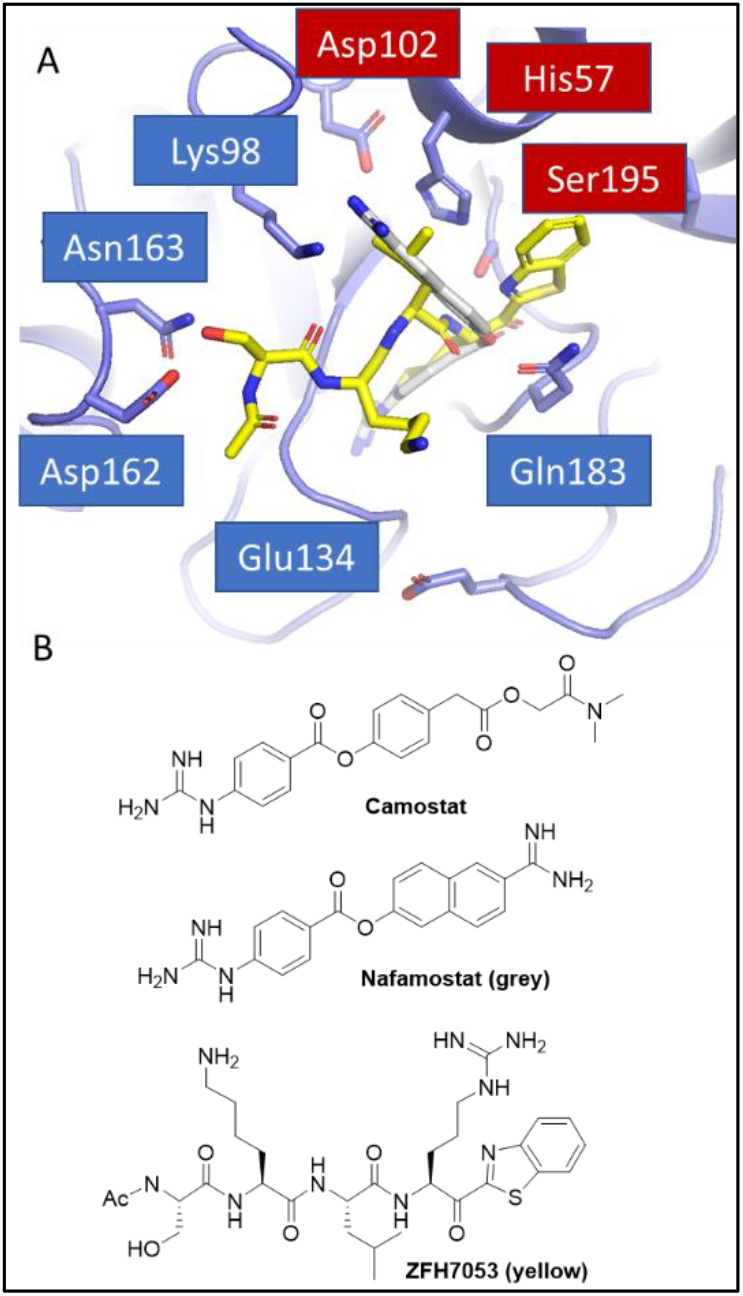
A) Docking model of compound **3** (yellow) and Nafamostat (grey) bound to a homology model of TMPRSS2 using Glides/Schrödinger; B) Structures of **3**, Camostat and Nafamostat.

#### Rational design of novel TMPRSS2-selective ketobenzothiazole (kbt) inhibitors

We have employed X-ray co-crystal structure data in the rational design of optimized HGFA, matriptase and hepsin inhibitors with increased potency and selectivity^*30–33*^. Shown in **Figure 3**, here we used a homology model of human TMPRSS2^*41*^ to computationally model our existing inhibitors (using Glide in Schrödinger) where we docked compound **3** (yellow; **Figure 3A**) and Nafamostat (grey). For Nafamostat, the naphthyl portion extends deep into the S2 pocket with the benzguanidine in the S1 pocket binding to the conserved S1 Asp189. The P4 Ser of **3** makes two H-bonds to Asp162 and Asn163 outside the S4 pocket, and the Ser backbone carbonyl is also within H-bonding distance to the P2 Lys98. The P3 Lys is near the S4 Glu134 making an electrostatic interaction while the P2 Leu resides in the S2 pocket and the benzothiazole fills the S1′ area. The reactive ester of Nafamostat and ketone of the kbt are adjacent to the Ser195-His57-Asp102 catalytic triad. Our model of **3** and Nafamostat bound to TMPRSS2 (**Figure 3A**) shows a nice fit to the S1-S4 and S1′ pockets of TMPRSS2. For **3**, it appears that the P4 Ser is making a dual H-bond to the Asp162 and Asn163 residues in TMPRSS2. This area is occupied by a Gln in both hepsin and matriptase but an Asp in HGFA and interestingly the additional Asn163 residue is unique to TMPRSS2. This suggests that a free amine in the P4 could provide selectivity over matriptase and hepsin. The structure also shows that Lys87 resides in the S2 pocket, suggesting a P2 Glu or Asp might be ideal for an inhibitor of TMPRSS2 and lead to potential selectivity over HGFA (P2 Ser), matriptase (P2 Phe) and hepsin (P2 Asn). This Lys87 is also near the P4 Ser sidechain indicating an Asp or Glu in this position could potentially make a third electrostatic interaction to Lys87. Further, the S2 pocket is large and appears that it could accommodate larger sidechains such as Phe, Tyr, or Trp.

#### PS-SCL Protease Substrate Specificity Profiling of TMPRSS2

To augment our rational design of more potent and selective TMPRSS2 inhibitors, we analyzed existing data on the peptide substrate preferences of TMPRSS2, HGFA, matriptase and hepsin. When comparing the positional scanning of substrate combinatorial libraries (PS-SCL) data of TMPRSS2 (**Figure 4**)^*25*^ with that of matriptase ^*42*^, hepsin ^*43*^ and HGFA^*31*^, it became apparent that there was significant overlap in their preferences for substrates. This data reveals TMPRSS2 is tolerant of many different P2 sidechains but prefers Phe and Ala/Thr like matriptase (**Figure 4**) which can also tolerate both large and small groups (prefers Ser/Ala) but hepsin and HGFA both prefer Leu. For P3 TMPRSS2 prefers Gln/Glu and Met whereas it is Lys/Gln for hepsin and matriptase and Lys/Arg for HGFA (**Figure 4**). The clearest distinction is in the P4 position where HGFA, matriptase and hepsin all prefer basic residues like Lys/Arg while TMPRSS2 prefers Ile/Gly and Pro, which is shared attribute with hepsin.

**Figure 4.**
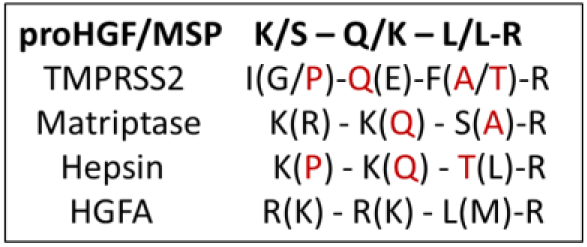
Combined PS-SCL for substrate specificity of TMPRSS2, matriptase, hepsin and HGFA. Using protease substrate terminology, the cleavage site for protease protein substrates is between the P1 and P1′ positions of the **P**eptide substrate, while S1, S1′ refers to the **S**ubsite ^*1*^ of the protease where the P1, P1′ amino acid sidechain binds.

#### MSP-MS profiling of TMPRSS2 for extended substrate sequence specificity

To strengthen our compound design, we obtained further information on the extended substrate specificity of TMPRSS2 using multiplex substrate profiling by mass spectrometry (MSP-MS).^*44*^ In the MSP-MS assay, a physiochemically diverse library of 228 tetra decapeptides was incubated for several hours with human recombinant TMPRSS2 in activity buffer. At different time points (15, 60 and 240 minutes) aliquots of the reaction mixture were extracted, quenched with 8M guanidium HCl and analyzed by tandem mass spectrometry to allow monitoring of protease-generated cleaved products. As a TTSP with a trypsin fold, TMPRSS2 is known for its high affinity towards substrates containing an arginine residue in the P1 position.^*25*^ This prominent P1 specificity was confirmed by our MSP-MS analysis, which showed a nearly exclusive occupancy of the P1 position by either Arg or Lys on TMPRSS2-generated cleaved products (**Figure 5A**). Peptide sequencing by LC-MS/MS enabled the identification of the 25 most preferred substrates for TMPRSS2 in our peptide library (**Figure 5B**). An IceLogo frequency plot (**Figure 5C**) displays the extended substrate specificity profile of TMPRSS2 at pH 7.5, which reveals the preference for hydrophobic amino acids in both P2 and P1′ positions, which flank the cleavage site. The analysis indicates a preferred cleavage of N-terminal tetrapeptides to be PLFR with other P4 amino acids H and M and P3 amino acids G, Y, V, and Q but only F at P2 which is striking. Largely this data recapitulates what was elucidated in the PS-SCL study (**Figure 4**). Based on careful analysis of the computational modeling work with these and the PS-SCL results, we selected additional existing compounds in our library of HGFA/matriptase hepsin inhibitors for testing and also synthesized four new analogs specifically designed for TMPRSS2: Ac-GQFR-kbt (**4**), Ac-PQFR-kbt (**5**), Ac-QFR-kbt (**6**) and Ac-IQFR-kbt (**7**).

**Figure 5.**
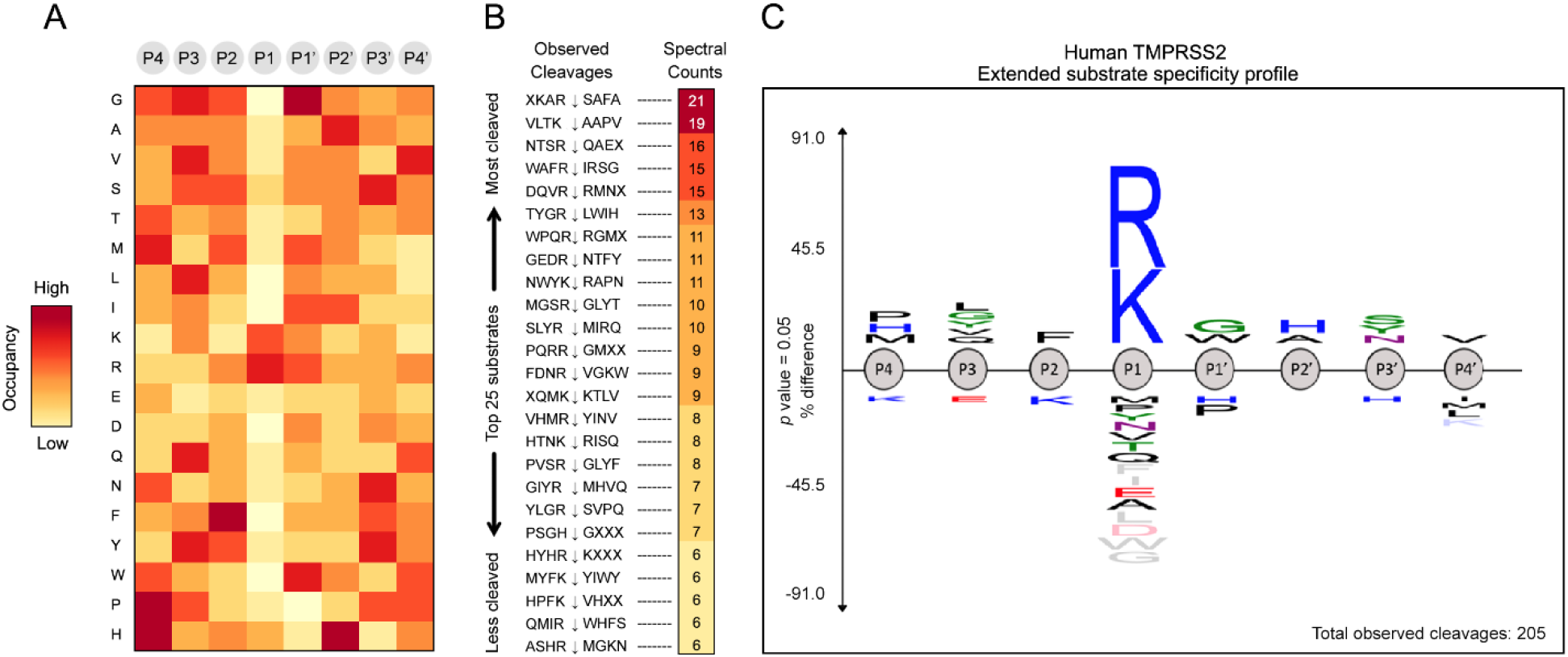
A) Heatmap displaying the overall amino acid frequencies in each of the P4-P4′ positions of a positive set of peptides cleaved by human TMPRSS2. Positive enrichment is observed as red and zero was set to yellow. B) Total spectral counts determined by LC-MS/MS highlight peptide sequences as high turnover rates substrates for TMPRSS2. C) IceLogo depicting the extended substrate specificity of human TMPRSS2 based on 205 cleavage events detected through MSP-MS analysis.

#### Inhibition of VSV-SARS-CoV-2 viral entry

Analogs based on the initial hit compounds **1** and **2** along with the substrate specificity data were Ac-SKLR kbt (**3**), Ac-WFR-kbt (**8**), Ac-SKFR-kt (**9**), Ac-KQFR-kt (**10**), Ac-SQLR-kt (**11**), Ac-FLFR-kbt (**12**), Ac-dWFR-kbt (**13**), dWFR-kbt (**14**), dWFR-kbt-CO2H (**15**), Ac-WLFR-kbt (**16**), Ac-KQLR-kbt (**17**), Ac-LLR-kt (**18**), Cyclo(DMK)R-kbt (**19**), Cyclo(DQK)R-kbt (**20**), and Cyclo(allylGLY)R-kbt (**21**). The majority of these compounds contain a Phe (F) residue in the P2 position as predicted from the PS-SCL study while others contain a Gln (Q) at the P3 position. When tested in the VSV pseudotype assay all displayed improved activity relative to **1** (**Table 1**) where the best compound was **13**. **13** is a tripeptide containing an unnatural amino acid D-Trp in the P3 position and Phe in the P2 position, predicted from PS-SCL and MSP-MS analysis to be preferred. **17** which contains a P3 Gln also predicted from PS-SCL shows the next best activity with an EC_50_ of 78 nM. Furthermore, **16** also having a P2 Phe showed excellent potency of 150 nM. Corresponding cycloamide analogs of **2**, **19** and **20** retained good activity (EC_50_ of 119 and 138 nM) for inhibition of viral entry of the VSV-pseudotype, but allyl phenyl ether analog **21** showed a significant decrease in activity (EC_50_ 565 nM).

**Table 1.**
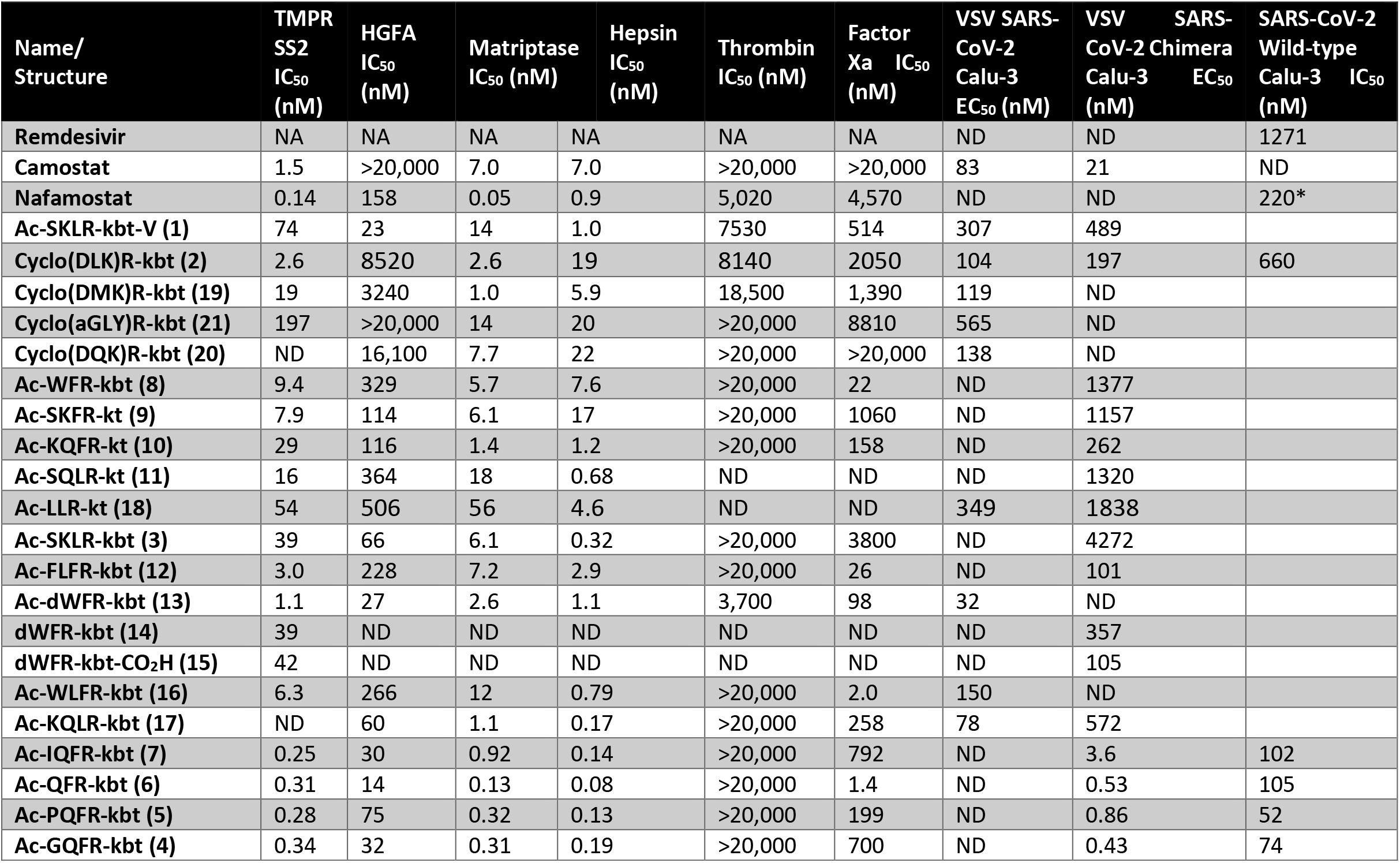
Structures and inhibition data of compounds for protease enzyme activity and VSV pseudotype and chimera SARS-CoV-2 viral entry into human Calu-3 lung epithelial cells (ND = not determined, NA = not applicable; *n=1).

The remaining compounds were tested in the VSV SARS CoV-2 chimeric assay. Excitingly, the most promising results (**Table 1**) were derived from our new rationally designed TMPRSS2 inhibitors, **4-7**. These compounds were designed by incorporating the preferred sidechains (as determined form PS-SCL data above) at all the P1-P4 positions of the inhibitor. The set of existing compounds all showed good potency with the best **12** showing an EC_50_ of 101 nM in blocking viral entry of VSV SARS-CoV-2 chimeras into Calu-3 cells, 6-fold more active than the second-best analog in the pseudotype assay, **17**. Excitingly, three of the new rationally designed analogs **4-6** displayed significantly increased potency (**Figure 6**) with sub nanomolar EC_50_s and the fourth new compound **7** is still extremely potent with an EC_50_ of 3.6 nM. This not only constitutes a 230-fold increase in activity for **4** (EC_50_ 0.43 nM) relative to the previous best compound **12** but also a 80-fold improvement over Camostat. Of the other compounds, similar potency compared to **12** was seen from tripeptide **15** (EC_50_ 105 nM) but less so from **14**, analogs of **13** which was the best compound tested in the pseudotype assay. Also showing excellent potency is **10** with an EC_50_ of 262 nM.

**Figure 6.**
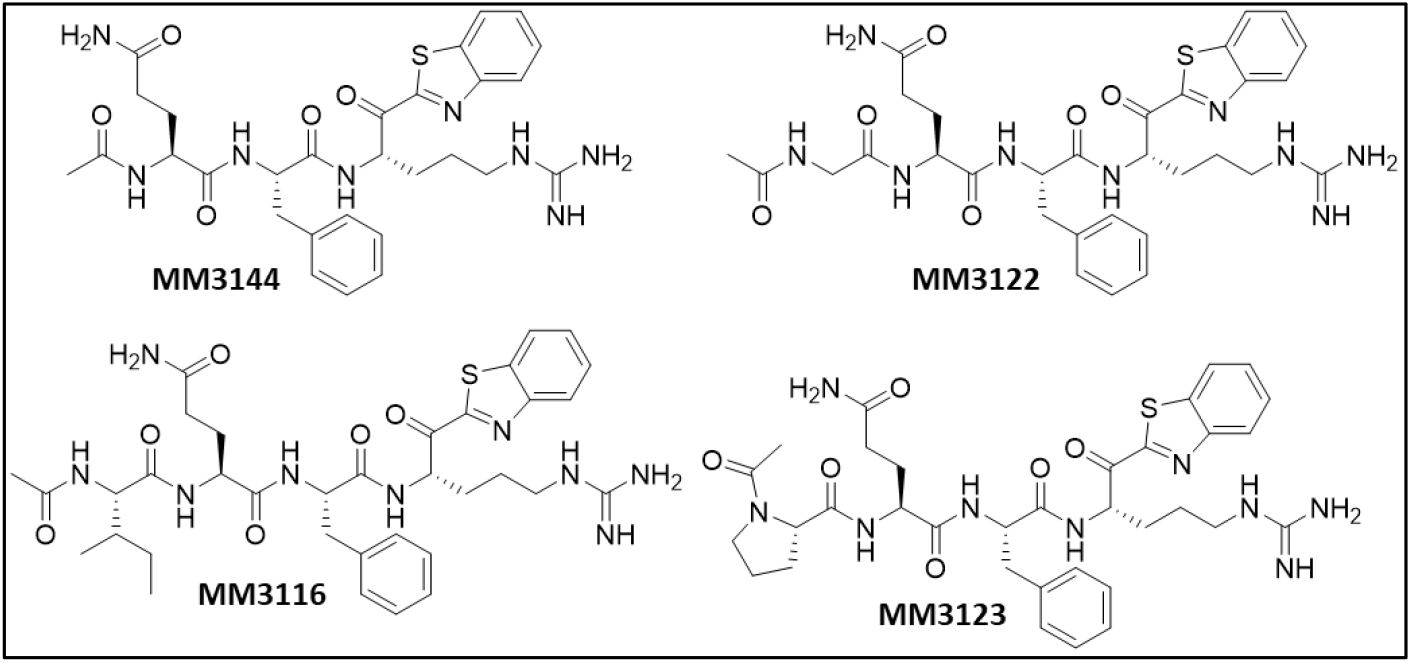
Rationally designed ketobenzothiazole (kbt) inhibitors of TMPRSS2.

#### Inhibition of VSV-MERS viral entry

To demonstrate broad coronavirus activity, we tested Camostat, **2** and our best new compounds, **4-7** (**Figure 6**) based on the VSV chimera results for their activity in blocking host cell entry of a MERS-CoV chimeric virus. Shown in **Table 2**, all compounds inhibited MERS-CoV viral entry with nearly equivalent potency as compared to SARS-CoV-2 (**Table 1**). Though the TMPRSS2-dependence of MERS-CoV has been demonstrated previously, this provides strong evidence that our compounds have broad-spectrum potential against other coronaviruses in the clinic. To confirm that this activity is indeed TMPRSS2-related, we next set out to produce recombinant TMPRSS2 protein for testing our compounds for their enzyme inhibitory activity.

**Table 2.**
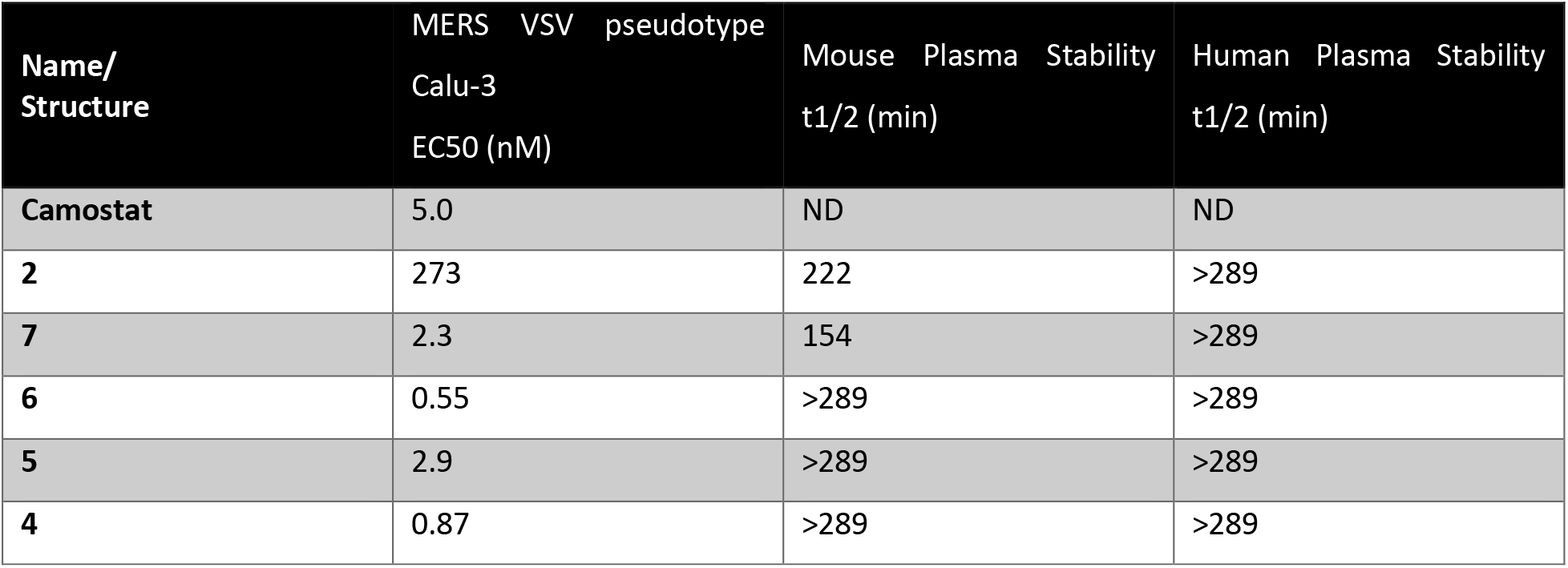
Activity of lead compounds against VSV pseudotype MERS viral entry into human Calu-3 lung epithelial cells. Also shown is the plasma stability of selected compounds in both mouse and human plasma (ND = not determined).

#### Expression and Purification of Recombinant TMPRSS2 (Protease Domain)

Recombinantly expressed TMPRSS2 protease domain (**Figure 7A**) was purified via Ni-NTA column (**Figure 7C**) then verified by SDS-PAGE and western blot (**Figure 7D** and **7E**) showing no presence of other proteins. Catalytic activity was tested by combining 150nM of purified TMPRSS2 protease domain with (MCA)-K-KARSAFA-K-(DnP), an 8-mer peptide FRET substrate that was synthesized based on the peptide that was preferred to be most efficiently by TMPRSS2 from our MSP-MS analysis (**Figure 5**). Michaelis-Menten kinetics showed a Km of 0.6±0.05 uM (**Figure 7B**). Relative to the full-length TMPRSS2 (see below), the k_cat_ value achieved for the protease domain (0.07/s) was 1000-fold lower than that observed for the construct bearing the LDLR class A domain and SRCR modules N-terminal to the catalytic domain (k_cat_ = 94,54/s). Further investigations are necessary to understand these differences and the interactions among the catalytic domain, the LDLR and the SRCR domains that impact the extended specificity and potentially the catalytic activity of TMPRSS2.

**Figure 7.**
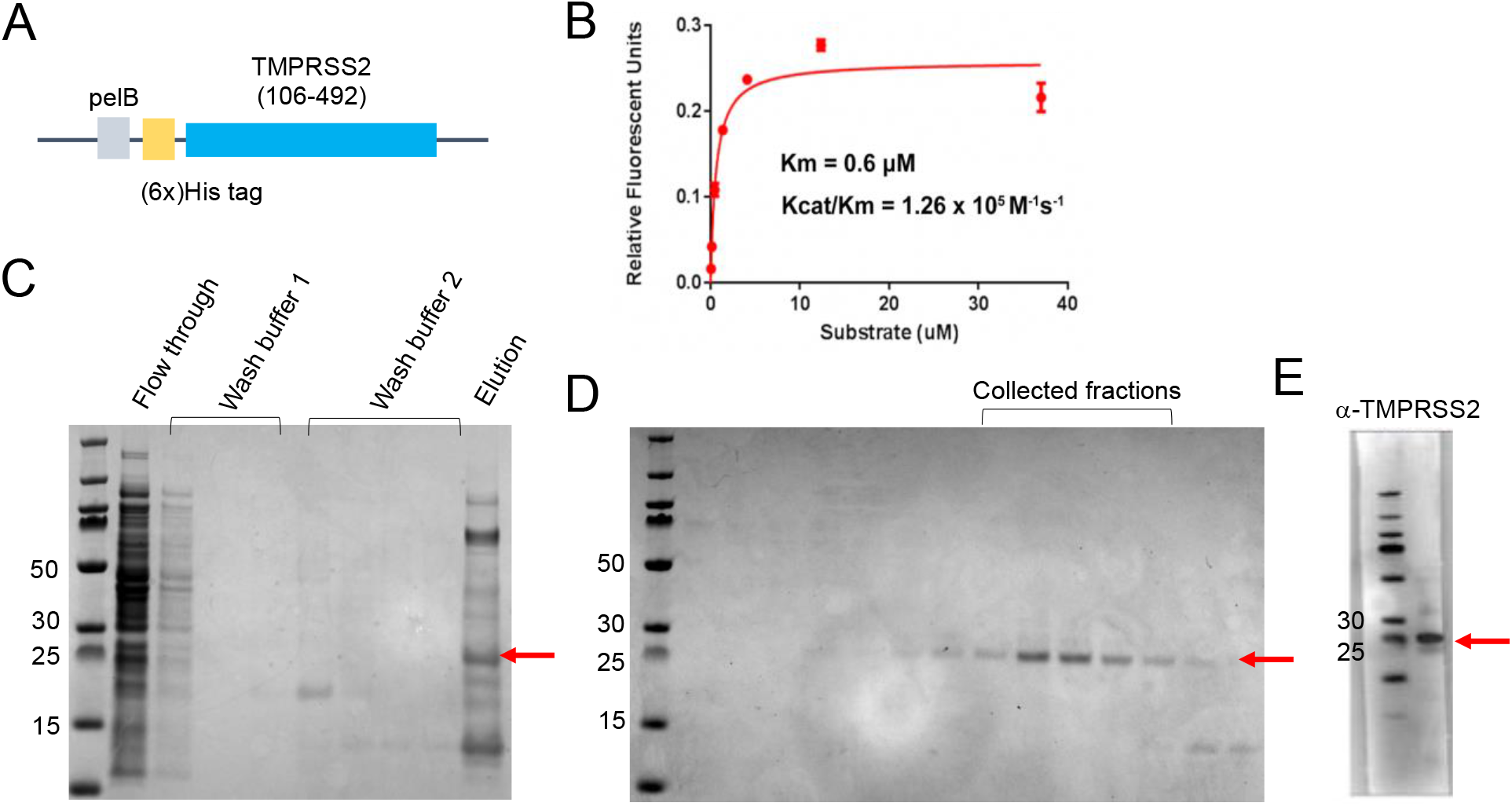
Bacterial periplamic expression and purification of TMPRSS2-protease domain. (A) TMPRSS2-protease domain expression construct. N terminal fused pelB signalling peptide allows for secretion of the target protease to the periplasm allowing for correct folding and ease in purification downstream. (B) Michaelis-Menten curve of TMPRSS2-Protease domain against (MCA)-K-KARSAFA-K-(DnP). Initial velocities of peptide cleavage were plotted against substrate concentration. Kinetic values were calculated using GraphPad Prism. (C) Denaturing SDS-PAGE gel of Ni-NTA affinity purification of TMPRSS2-Protase domain. Correct band size for TMPRSS2-protease domain (~26kDa) shown in Red arrow. (D) SDS-PAGE gel of size exclusion chromatography fractions. Fractions only containing protease domain were collected and pooled. E) Western blot against pooled TMPRSS2-protease domain. Blot using monoclonal TMPRSS2-protease domain antibody (M05), clone 2F4 confirms purity of sample and no existence of degradation products.

#### Recombinant TMPRSS2 (Full-length) Enzyme Assay

Using active recombinant full-length TMPRSS2 as described above, along with Boc-QAR-AMC as a fluorogenic substrate^*40*^, we found the K_m_ was 85.6 μM using an enzyme concentration of 3 nM (Supplementary material). We used this substrate to test all inhibitors for inhibition of TMPRSS2 proteolytic enzyme activity in a standard kinetic assay using Nafamostat and Camostat as controls, where we determined the compound IC_50_s over the period of one hour, following 30 min compound incubation with enzyme. The IC_50_ values for Camostat and Nafamostat are 1.5 nM and 0.14 nM which is similar but slightly improved compared to values previously reported of 6.2 nM and 0.27 nM, respectively.^*40*^ We found the IC_50_ data we generated for TMPRSS2 inhibition closely correlated with the VSV pseudotype and chimera assay data with some exceptions. Multiple compounds were significantly more potent than Camostat and several were equipotent to Nafamostat (**Table 1**). The new rationally designed lead compounds **4-7** (**Figure 6**) all displayed exquisitely potent sub nanomolar IC_50_ values of which **7** was the best with an IC_50_ of 250 picomolar similar to that of Nafamostat. It should be noted however that Nafamostat is an irreversible inhibitor while the kbt class of inhibitors are reversible, so it is difficult to directly compare IC_50_ values between the two series having different mechanism of inhibition. While the initial hit compound **1** still shows good potency against TMPRSS2 (IC_50_ 74 nM), it was significantly weaker when compared to the rationally designed TMPRSS2 inhibitors which was expected. However, the other initial hit compound **2** is a much more potent inhibitor of TMPRSS2 activity than **1** with an IC_50_ of 2.6 nM, only about 10-fold less active than **4-7**. It is noteworthy that the P3-P1 tripeptide **6** without the P4 sidechain was almost as active as either of the tetrapeptides **4**, **5** or **7** which have a P4 residue. However, this compound **6** and the other tripeptides including **8** and **13** suffer from high inhibition of the plasma protease Factor Xa which is undesirable for further drug discovery due to potential bleeding side effects in patients.

The least active compound was the aryl ether cyclic peptide **21** with an IC_50_ of 197 nM which is however still respectable. It is presumed the larger, more constrained aryl ether ring system is not an ideal fit for binding to the TMPRSS2 active site through bridging the S2-S4 pockets. It is important to note that while TMPRSS2 activity is potent in all inhibitors, most compounds tested are all still relatively active against the other proteases HGFA, matriptase and hepsin.

The most selective TMPRSS2 analogs are **16** with an IC_50_ of 6.3 nM, being 40-fold more active than HGFA, 2-fold over matriptase and **12** which is 60-fold more selective over HGFA and 2-fold against matriptase. As with other compounds in this series of peptidyl kbt inhibitors, in our experience we have found that deriving selectivity over hepsin is challenging but one of our best examples turns out to be initial hit **2**, which is 4-fold more active against TMPRSS2 relative to hepsin and almost 10-fold more active against matriptase. The consequences of inhibiting these other serine proteases, especially the other HGF-activating proteases, in the COVID-19 scenario is unclear at this time but may be beneficial for treatment rather than detrimental.

There are clear structure-activity relationships (SAR) from the 21 compounds tested. For example, it appears that TMPRSS2 prefers large groups extending beyond the kbt S1′ C-terminal portion of the inhibitor since **1** have increased activity relative to unsubstituted kbt analog **3**.

Strengthening this hypothesis, the compounds with the smaller ketothiazole (kt) warhead in the P1′ position like seen in **9-11** also lose potency relative to analogs with the larger kbt. Also, the P3 position is important for activity where it seems basic groups like Lys (K) are undesired (**9**, **3**, **1**) which seems also true for the P4 position as seen with compound **17**. Finally, as predicted from modeling and the PS-SCL data, Phe is clearly preferred in the P2 position. Therefore, it is likely that diverse new analogs can be pursued with other aromatic sidechains like Trp and Tyr which will also produce inhibitors with exquisite TMPRSS2 activity and potentially more selectivity.

Given the homology model of TMPRSS2 used in this study was exclusive to the catalytic domain of TMPRSS2, both catalytic domain and full length TMPRSS2 were tested against candidate compounds. The inhibition IC_50_ values against the protease domain versus full length TMPRSS2 shows strikingly different values with estimated IC_50_s being over 100 μM compared to picomolar to double digit nanomolar IC_50_ values for the commercial full length TMPRSS2 (**Table 1** and Supplementary Data). It is not clear from our current studies what the direct cause of this noticeable difference is, but we suspect that the protein domains/modules of TMPRSS2 which are N-terminal to the catalytic domain play an important role in ligand interactions and subsequent catalysis.

#### Antiviral activity of lead compounds (Wild-type SARS-CoV-2)

After confirming and quantitating TMPRSS2 enzyme inhibition of all compounds against recombinant protein and their selectivity over HGFA, matriptase, hepsin, thrombin, and Factor Xa, we assessed the activity of our most promising lead compounds **2**, **4-7** towards wild-type SARS-CoV-2 by employing a CellTiter-Glo (Promega) cellular viability assay using lung epithelial Calu-3 cells.^*45*^ We used Remdesivir and Nafamostat (n=1) as a positive control. While all compounds showed excellent activity in this assay (**Figure 8** and **Table 1**), the most potent compounds were the rationally designed TMPRSS2 inhibitors **4** and **5** (**Figure 6**) with EC_50_s of 74 and 52 nM, respectively which is >20-fold improved over Remdesivir and also significantly more active than Nafamostat (note: based on n=1). Importantly, all five kbt inhibitors showed no signs of toxicity to Calu-3 cells (up to 50 μM, which is nearly 1000 times the IC_50_ for **5**, while Remdesivir (and Nafamostat) displayed toxicity at the highest concentration tested (50 μM).

**Figure 8.**
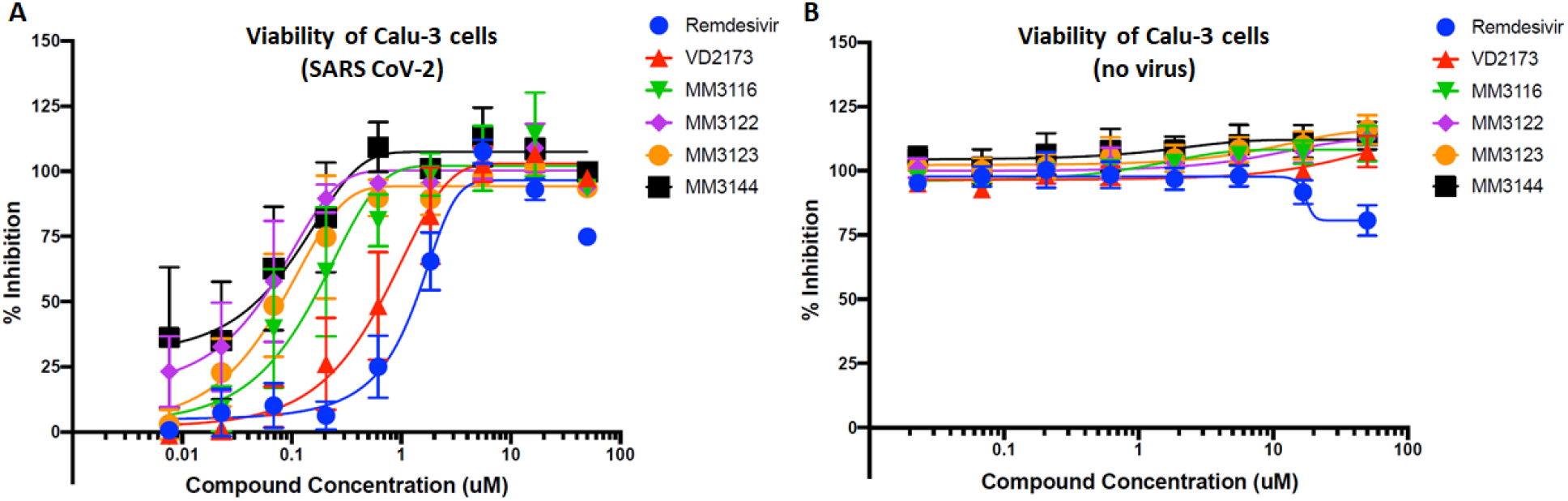
A) Inhibition by lead TMPRSS2 inhibitors of Calu-3 viral toxicity of wild-type SARS-CoV-2 using the CellTiter-Glo (Promega) assay. B) Viability of Calu-3 cells with compounds in the absence of virus.

#### Pharmacokinetics and metabolic stability of VD2173 (2) and MM3122 (4)

VD2173 (**2**) and lead compounds **4-7** were tested for their in vitro stability in mouse and human blood plasma (**Table 2**). All compounds have excellent stability in both mouse and human plasma with >289 min half-life except for **7** which had a half-life of 154 min in mouse plasma and 2 with a 222 min half-life. Based on potency, selectivity and in vitro properties, compound **4** (**MM3122**) was selected as a lead candidate and was tested for its in vivo pharmacokinetics (PK) in mice. The mouse PK of VD2173 in plasma has been published elsewhere.^*36*^ Shown in **Figure 9**, VD2173 has an outstanding half-life of 8.7 h in mice with a high AUC and exposure in plasma beyond 24 h. Excitingly, rationally designed TMPRSS2 lead compound MM3122 also had excellent PK with a half-life of 8.6h. Compounds **5-7** were not tested for PK due their lower antiviral activity and the relatively high Factor Xa activity of **5** and **6**. Both VD2173 and MM3122 were subsequently tested for their lung exposure over a period of 24 h after IP dosing. It was found that both compounds attained high levels of compound in the lung and had excellent AUC but with MM3122 being superior with a 7.5 h half-life versus 4.2 h for VD2173. The potency of MM3122 versus VD2173 in both the TMPRSS2 enzyme assay and viral assays is magnitudes higher (500-fold) making MM3122 an ideal lead candidate drug suitable for in vivo efficacy studies of COVID-19 and further optimization to be reported in a future communication.

**Figure 9.**
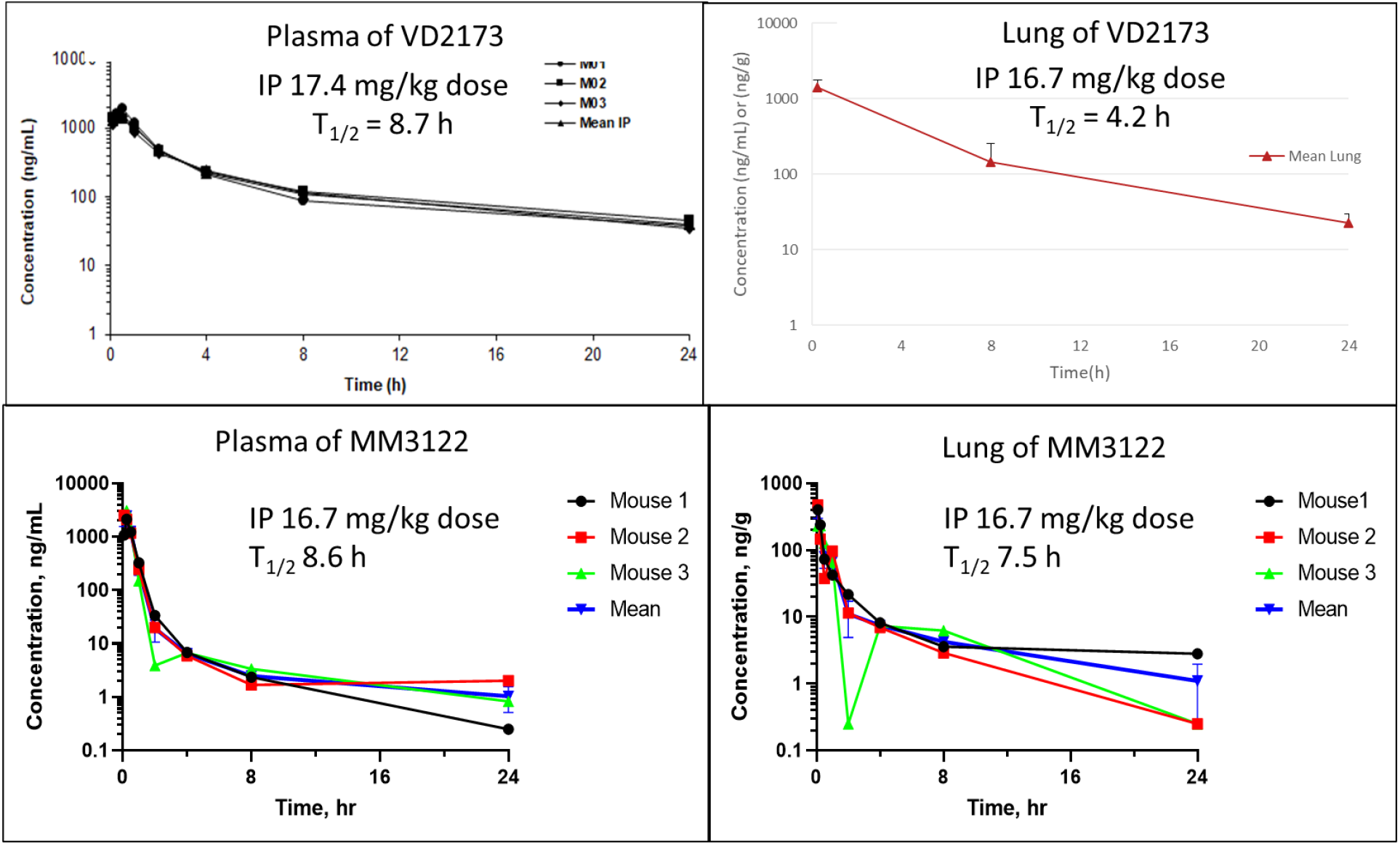
Mouse pharmacokinetics of VD2173 (1) and MM3122 (4) (IP nominal dose 20 mg/kg). Concentrations of compound over time in plasma and lung are shown.

#### Acute 7-day toxicity study of MM3122 (4) in mice

We further tested lead compound MM3122 for its safety in mice. MM3122 was administered daily to mice at three different single dose levels of 20, 50, and 100 mg/kg via intraperitoneal (IP) injection over a period of 7 days. No adverse effects were observed in any of the treatment groups and no weight loss or changes to harvested organs (liver, spleen, and kidneys) were noted compared to the control group (Supplementary material).

#### Conclusions

TMPRSS2 has been shown to be essential for host cell viral entry and replication of SARS-CoV-2. Based on molecular docking studies using a published homology model of TMPRSS2^*41*^ and substrate specificity data from PS-SCL, we hypothesized that a set of our existing peptidyl ketobenzothiazole inhibitors of HGFA, matriptase and hepsin would also inhibit TMPRSS2. Indeed, we demonstrated that these compounds not only potently inhibit TMPRSS2 enzyme activity of recombinant and cell-surface protein, but also host cell entry and toxicity driven by the Spike protein of SARS-CoV-2 into ACE2/TMPRSS2-expressing Calu-3 lung epithelial cells. After further optimization, we identified multiple potent inhibitors of TMPRSS2, with four analogs rationally designed for TMPRSS2 displaying sub nanomolar activity in both the enzyme assay and in blocking the entry of VSV-SARS-CoV-2 chimeras into human Clau-3 epithelial lung cells. In addition, several of these compounds displayed excellent potency against MERS, another prominent coronavirus. We further confirmed potent antiviral activity against the wild-type SARS CoV-2 virus with five lead compounds thus identifying the most promising lead compound MM3122 (**4**). We have clearly established that this novel TMPRSS2 inhibitor MM3122 is more potent than Remdesivir, Camostat and Nafamostat. Importantly, this compound also shows no toxicity to Calu-3 cells in contrast to Remdesivir and Nafamostat which are toxic at higher concentrations. MM3122 has excellent metabolic stability in both mouse and human plasma as well as outstanding pharmacokinetics and safety in mice. Cyclic peptide VD2173 (**2**) and rationally designed TMPRSS2 inhibitor MM3122 both have an impressive >8 h half-life in mice. In addition to its excellent TMPRSS2 inhibition, pharmacokinetics, and antiviral activity, VD2173 has anticancer efficacy in animal models of lung cancer.^*36*^ Judging from the protease selectivity data and potency of MM3122 and other new compounds, these are predicted to also show corresponding anticancer activity, in particular for HGF-driven prostate cancer for which TMPRSS2 is known to play a key role^*25, 46*^.

MM3122 represents an advanced lead candidate for clinical development as a novel antiviral drug for COVID-19 and against infections caused by other coronaviruses like MERS in which we showed equipotent activity of MM3122. We are currently optimizing MM3122 and this class of inhibitors for TMPRSS2 selectivity and antiviral potency, and in due course will test MM3122 and other optimized leads in appropriate animal models of COVID-19. In addition to being novel drugs, selective TMRSS2 inhibitors can be used as valuable chemical probes to help elucidate mechanisms of viral pathogenesis, including host cell-virus interactions, Spike protein processing and ACE2 receptor binding. Since TMPRSS2 plays a key role as a viral protein processing protease in the pathogenesis of other coronaviruses (SARS-CoV, MERS-CoV) as well as influenza viruses ^*47–52*^, MM3122 and this new class of inhibitors may be effective not only against COVID-19 but for infections caused by most or all corona and influenza viruses. Thus, these small-molecule inhibitors of TMPRSS2 not only hold much promise as new drugs to treat SARS-CoV-2 infections but also potentially represent broad-spectrum antivirals.

## Experimental

### TMPRSS2-Protease domain expression and purification

Periplasm secreted bacterial expression of serine proteases has been widely reported to yield correctly folded high-quality enzyme due to its reducing environment allowing for disulfide formation and fewer proteases being present compared to the cytoplasm.^*53–54*^ We therefore employed a similar strategy for TMPRSS2 protease domain. The human TMPRSS2 extracellular domain (residues 106-492) was cloned into a pET28a vector with a *pelB* leader sequence followed by an N-terminal 6x His tag (**Figure 7A**) using standard Gibson assembly procedures.^*55*^ E. Coli DH5a cells were transformed with the Gibson assembly reaction mixture and resulting plated colonies were selected and verified with Sanger DNA sequencing for correct assembly of the cloned product. Correctly cloned plasmid was used for transforming *E. Coli* BL21(DE3) cells and later used for subsequent expressions. A single colony of BL21(DE3) cells was picked and inoculated into 50 ml of LB 2% glucose, 50 mg/ml Kanamycin (Km) culture and grown overnight at 37 °C 220 rpm. Liter scale liquid media (LB 0.1% glucose, 50 mg Km) were inoculated with overnight cultures at a starting OD600 of 0.05 and expression was induced with a final concentration of IPTG at 1mM at approximately OD600 of 0.7. Expressions were carried out at 16 °C for approximately 72 hours. Cells were collected by centrifugation and the periplasmic extracts were prepared by osmotic shock. Briefly, cell pellets were resuspended in TES buffer (200 mM Tris-HCl pH 8, 0.5 mM EDTA, 500 mM sucrose) and incubation at 4 °C for 1 hour followed by addition of cold water and incubation at 4 °C for 45 minutes. The periplasmic extract was isolated with centrifugation (10,000 g for 30 min) at 4 °C and imidazole (10 mM final concentration) and MgCl_2_ (100 ml per liter expression culture volume) was added for overnight batch binding with Ni-NTA resin (2 ml slurry per liter of expression culture volume). Purification was carried out with 10 column volumes of Wash buffer 1 (50 mM Tris 250 mM NaCl pH 7.6), subsequently 10 column volumes of wash buffer 2 (50 mM Tris pH 7.6, 250 mM NaCl, 20 mM imidazole) and eluted with elution buffer (50 mM Tris pH 7.6, 250 mM NaCl, 4 mM benzamidine, 1 mM CaCl_2_, 500 mM imidazole) (**Figure 7C**). SDS-PAGE was performed with fractions of eluate, and those showing the correctly sized band for TMPRSS2 protease domain were pooled. Consolidated fractions were buffer exchanged against 25 mM Citrate pH 6, 250 mM NaCl, 4 mM benzamidine, 1 mM CaCl_2_ to remove excess imidazole and prevent autolysis and concentrated down in the same buffer with 10kDa NMWL Amicon ultra filtration units. Potential aggregates and degradation products were removed by size exclusion chromatography with 25 mM Citrate pH 6, 250 mM NaCl, 4 mM benzamidine, 1 mM CaCl_2_ and 10% glycerol and Superdex 200 Increase 10/300 GL. TMPRSS2 protease domain containing fractions by SDS-PAGE (**Figure 7D**) were pooled. A final western blot against the protease domain (TMPRSS2 monoclonal antibody (M05), clone 2F4) was performed to verify collected sample to be TMPRSS2-protease domain. Pooled sample was flash frozen for subsequent assays (**Figure 7E**).

### Recombinant enzyme inhibition assays

Recombinant human TMPRSS2 was purchased from Cusabio Technology (CSB-YP023924HU) and assayed in 25 mM Tris-HCl, 150 mM NaCl, 5 mM CaCl_2_, 0.01% Triton X-100, pH 8.0 at a final concentration of 3 nM. To calculate K_M_, Boc-QAR-AMC (Vivitude, MQR-3135-v) was serially diluted in DMSO and then each diluent was further diluted in assay buffer such the final substrate concentration in the assay ranged from 0.514 μM to 200 μM with a final DMSO concentration of 0.5%. Assays were performed in a total volume of 30 uL in triplicate wells of a black 384-well plates (Nunc 262260) and initial velocity was measured in 35 second intervals for 7 minutes with excitation of 360 nm and emission of 460 nm on a Biotek HTX platereader. K_M_ and V_max_ were calculated from Michaelis-Menten plots using GraphPad Prism. For inhibitor studies, compounds were serially diluted 3-fold in DMSO and then preincubated with TMPRSS2 in assay buffer for 30 min at room temperature. The reaction was initiated following the addition of Boc-QAR-AMC and fluorescence was measured in 190 second intervals for 90 min. The final concentration of enzyme and substrate were 3 nM and 86.6 μM, respectively and the inhibitor concentrations ranged from 2 μM to 0.15 pM. Assays were performed in quadruplicate plates and IC_50_ was calculated from dose-response curves using Graphpad Prism. IC_50_ values for HGFA, matriptase, hepsin, factor Xa and thrombin were determined using our published assays^*32–34*^.

### Multiplex Substrate Profiling by Mass Spectrometry (MSP-MS)

Human recombinant TMPRSS2 (100 nM) expressed in yeast* was incubated at room temperature with a physiochemically diverse library of 228 tetradecapeptides to a final concentration of 500 nM. Aliquots were removed at different time intervals (15, 60 and 240 minutes) and subsequently quenched with 1Eq volume of 8M guanidinium hydrochloride. Samples were desalted using C18 tips (Rainin) and analyzed by LC-MS/MS peptide sequencing using a Quadrupole Orbitrap mass spectrometer (LTQ Orbitrap) coupled to a 10,000 psi nanoACQUITY Ultra Performance Liquid Chromatography (UPLC) System (Waters) for peptide separation by reverse phase liquid chromatography (RPLC). Peptides were separated over a Thermo ES901 C18 column (75-μm inner diameter, 50-cm length) coupled to an EASY-SprayTM ion source and eluted by applying a flow rate of 300 nL/min in a 65-minute linear gradient from 2–50% in Buffer B (acetonitrile, 0.5% formic acid). Survey scans were recorded over a 325–1500 m/z range and up to the three most intense precursor ions (MS1 features of charge ≥ 2) were selected for collision-induced dissociation (CID). Data was acquired using Xcalibur software and processed as previously described.^*56*^ Briefly, raw mass spectrometry data was processed to generate peak lists using MSConvert. Peak lists were then searched in Protein Prospector v.6.2.2^*57*^ against a database containing the sequences from the 228 tetradecapeptide library. Searches used a mass accuracy tolerance of 20 ppm for precursor ions and 0.8 Da for fragment ions. Variable modifications included N-terminal pyroglutamate conversion from glutamine or glutamate and oxidation of tryptophan, proline, and tyrosine. Searches were subsequently processed using the MSP-xtractor software (http://www.craiklab.ucsf.edu/extractor.html), which extracts the peptide cleavage site and spectral counts of the corresponding cleavage products. Spectral counts were used for the relative quantification of peptide cleavage products.

### *Expression of Human Recombinant TMPRSS2 in *Pichia pastoris*

Human Recombinant TMPRSS2 was expressed and purified from yeast following protocols previously described^*25*^, with slight modifications. Briefly, the scavenger receptor cysteine-rich (SRCR) and serine protease domain regions of TMPRSS2-N249G were cloned into a pPICZα-B construct (EasySelect Pichia Expression Kit – Invitrogen) according to the manufacturer, transformed into *Pichia pastoris* strain X33, and selected on plates containing increasing concentrations of Zeocin. A single colony was grown in 10 mL buffered medium with glycerol (BMGY) overnight at 30°C and 230 rpm. The overnight culture was used to inoculate 1 L BMGY and grown until OD 2-6. Cells were pelleted and resuspended in 100 ml buffered medium with methanol (BMMY). Cells were induced with 5% methanol every 24 hours for 72-96 hours. Secreted TMPRSS2 was precipitated with 70% ammonium sulfate at 4°C overnight, pelleted at 27,000 x g for 45 minutes and resuspended in 50 mM Tris pH 8, 0.5 M NaCl, 0.01% CHAPS, and solubilized for 2 hours at 4°C. The solubilized protein was then purified over a gravity column containing soybean trypsin inhibitor immobilized agarose (Pierce). TMPRSS2 was concentrated and then purified with size exclusion chromatography using 50 mM potassium 83 phosphate buffer pH 6, 150 mM NaCl, 1% glycerol to prevent auto-proteolysis.”

### Cell-based TMPRSS2 fluorogenic enzyme inhibition assay

A PLX304 plasmid–containing human TMPRSS2 open reading frame from the ORFeome Collaboration (Dana-Farber Cancer Institute, Broad Institute of Harvard and Massachusetts Institute of Technology [HsCD00435929]) was obtained from DNASU Plasmid Repository, and a control PLX304 vector was obtained from Addgene (Watertown, MA, USA). HEK-293T cells were grown in DMEM supplemented with 10% FBS and seeded in a black, 96-well plate (75,000 cells/well). The following day, cells were transfected overnight with either a control plasmid (PLX) or TMPRSS2 (PLX-TMPRSS2) via TransIT LT-1 transfection reagent (Mirus Bio) in 100 μL of OptiMEM per well. Twenty-four hours after transfection, the media was replaced with 80 μL of phosphate-buffered saline (PBS). Inhibitors or PBS alone were added to the wells in the indicated 5 concentrations and incubated at 25°C for 15 minutes. The fluorogenic substrate Boc-QAR-AMC (R&D Biosystems) was then added to each well to a final concentration of 100 μM. Fluorescence (excitation 365 nm, emission 410 nm) was kinetically measured every 15 minutes for a total time of 150 minutes at 37°C using a GloMax plate reader (Promega).

### Antiviral activity (VSV SARS-CoV-2 and MERS chimeras)

#### Viruses used

VSV-eGFP, a recombinant VSV expressing a GFP reporter (depends on the VSV glycoprotein G for entry), has been previously described.^*53*^ VSV-SARS-CoV-2, a replication competent infectious VSV chimera which employs the SARS-CoV-2 Spike (S) protein for viral entry in place of VSV G and expresses eGFP, has also been previously described.^*37, 59*^ VSV-MERS was created in the same manner as VSV-SARS-CoV-2, except the MERS Spike missing the terminal 21 amino acids (HCoV-EMC/2012 strain) was inserted in place of VSV G.

#### Experimental procedure

Human Calu-3 lung epithelial cells or Vero cells (African green monkey kidney) were seeded in a 96-well black plate in DMEM containing 10% FBS for 24 hours (37°C and 5% CO2). The next day, cells were pretreated for 2 hours with inhibitor or vehicle control (DMSO) in 50 μl serum-free DMEM, and subsequently infected with VSV-SARS-CoV2, VSV-MERS, or VSV-eGFP at a multiplicity of infection (MOI) of 0.5. At 7 hours post infection (single round of infection), cells were fixed in 2% formaldehyde and nuclei stained with 10 μg/ml Hoechst 33342 (Invitrogen) for 30 minutes at room temperature. Cells were washed once and stored in PBS after fixation, and automated microscopy performed using an InCell 2000 Analyzer (GE Healthcare) in the DAPI and FITC channels (10X objective, 9 fields per well, covering the entire well). Images were analyzed using the Multi Target Analysis Module of the InCell Analyzer 1000 Workstation Software (GE Healthcare) to identify the percentage of GFP-positive cells in each well (top-hat segmentation). The percentage GFP-positive cells for experimental conditions was normalized against the control (DMSO-treated) cells and expressed as relative infectivity. GraphPad Prism (version 8.4.2) was used to calculate the EC_50_ of each drug. Statistics (comparison of VSV-SARS-2 to VSV-eGFP with each drug; Student’s t-test) were performed in Microsoft Excel. Three biological replicates were performed.

#### Antiviral activity (VSV pseudotype SARS-CoV-2)

We analyzed pseudotype entry driven by the spike protein of SARS-CoV-2 (SARS-2-S) or the glycoprotein of vesicular stomatitis virus (VSV-G) into the TMPRSS2-positive human lung cell line Calu-3^*10, 24*^. VSV-G was used as a control, as it does not depend on TMPRSS2 for host cell entry. Besides Calu-3 cells, we further used Vero cells (African green monkey, kidney) as a control, as these cells do not express TMPRSS2 and therefore any reduction in SARS-2-S-driven entry would be either related to unspecific side effects or cytotoxicity. Each compound was tested in three separate experiments with independent pseudotype batches. We treated cells (96-well format) with different concentration of inhibitor or solvent (DMSO) diluted in medium (50 μl/well) for 2 h at 37 °C and 5 % CO2, then we added 50 μl of pseudotype on top and incubated for 16 h at 37 °C and 5 % CO2. Next, we measured virus-encoded firefly luciferase activity in cell lysates (indicator of pseudotype entry into target cells). The data was normalized against control (DMSO-treated cells = 100 % pseudotype entry) and plotted with GraphPad Prism (version 8.3.0), to calculate effective concentration EC50 and perform statistics (comparison against the respective control; two-way ANOVA with Dunnett’s posttest).

### Antiviral activity (Wild-type SARS-CoV-2)

We seeded Calu-3 cells onto a white, flat bottom 96-well plate at 10,000 cells per well in Dulbecco’s Modified Eagle Medium supplemented with 10% heat-inactivated fetal bovine serum, 100 U/mL Penicillin-Streptomycin (Life Technologies, Inc., 15140-163), and buffered with 10 mM HEPES and incubated in a humidified 5% CO_2_ incubator at 37°C. 16 hours later, the media was removed and replaced with the 50 μl/well of the same media but containing 2% heat-inactivated fetal bovine serum (D-2) instead of 10%. In D-2, compounds were serially diluted 3-fold in a 9-point series and added in a 25 ul volume to the 96-well plate to make final concentrations ranging from 50 μM to 7.6 nM on the assay plate for one hour. We then infected the cells (with compounds still on them) with 25 ul/well of SARS-CoV-2 at an MOI of 0.1 PFU/cell, with a final total volume of 100 μl/well. We incubated the plates for 72 hours as described above and then added 25 μL/well of CellTiter-Glo^®^ Reagent (prepared as directed by manufacturer, Promega, G7573), followed by 5 min shaking and 20 min room temperature incubation before detecting luminescence with a Biotek Synergy H1 plate reader. Compound cytotoxicity was determined using the same assay, but instead of adding virus, we added 25 μl of D-2. These experiments were performed three times, and we performed nonlinear regression on log(inhibitor) vs response with variable slope and four parameters on compiled results from the three experiments. Likewise, IC_50_s were calculated based on all three biological replicates.

### Animal studies

All animal procedures were performed following the guidelines and approval of the Washington University’s Institutional Animal Care and Use Committee. Animals were maintained in a controlled temperature (22°C± 0.5°C) and lighting (12L:12D) environment. Standard laboratory chow and water were supplied ad libitum.

### Mouse toxicity studies

NSG male and female mice (3 per group) were injected with 0, 20, 50, or 100 mg/kg MM3122 (in 1-5%DMSO/95% saline; IP) daily for 7 days. Animal health was assessed daily for drug toxicity (ruffled fur, discharge from the eyes, weight loss, dehydration, lethargy, hypothermia, abnormal tissue growth) and body weights were measured every 2-3 days. At the end of the experiment animals were sacrificed and organs inspected. Liver, spleen, and both kidneys were collected and weighed.

### Mouse pharmacokinetic (PK) studies

MM3122 was tested for its PK in mice (single IP dosing, actual dose 16.7 mg/kg). Eight animal groups contained three animals each (male, CD-1) with plasma drawn and lung tissue harvested at eight different sampling time points (0.08, 0.25, 0.5, 1, 2, 4, 8, 24 h) post dosing. VD2173 was tested for its PK in mice (single IP dosing, actual dose 16.7 mg/kg). Three animal groups contained three animals each (male, CD-1) with plasma drawn and lung tissue harvested from each group of animals at three different sampling time points (0.25, 8, 24 h) post dosing. Bioanalysis of plasma and lung tissue extracts were performed using standard LC/MS/MS techniques.

## Acknowledgements

We thank David Griggs and Scott Campbell at Saint Louis University for conducting the pharmacokinetic studies on MM3122. We also thank Zhenfu Han for previously synthesizing several of the compounds in this manuscript, but which have already been published elsewhere. We acknowledge Michael Winter at UCSF for his help with the initial MSP-MS and PS-SCL profiling. We would like to thank the Siteman Cancer Center (#16-FY18-02 and SCC P30CA091842) of Washington University and Barnes Jewish Hospital Foundation (BJHF 4984) in Saint Louis for funding (J.WJ.). This work was also funded by the following awards from the National Institutes of Health (NIH): R43 CA243941 (J.W.J. and L.K.), R43 CA224832 (J.W.J. and L.K.), U19 AI142784 (C.L.S.), P50AI150476 (C.S.C.), U19 AI070235 (M.E.R) and the Campaign Urging Research for Eosinophilic Diseases (CURED) Foundation (M.E.R.). Additional support was provided by a Fast Grant from Emergent Ventures at the Mercatus 9 Center, George Mason University (C.S.C.). S.P. acknowledges funding by BMBF (RAPID Consortium, 01KI1723D and 01KI2006D; RENACO, 01KI20328A; SARS_S1S2 01KI20396; COVIM consortium, 01KX2021), the country of Lower Saxony (14-76103-184) and the German Research Foundation (DFG) (PO 716/11-1, PO 716/14-1). Work with live SARS-CoV-2 was funded by a Burroughs Wellcome Fund Investigators in the Pathogenesis of Infectious Disease Award (C.L.S.).

## Supplementary Data

1. Single point inhibition data using recombinant protease domain/FRET substrate for **4-6**.
2. Protease selectivity data for **1** and **2** and Camostat/Nafamostsat.
3. Synthesis and NMR and HPLC-MS spectra of new compounds **2**, **4-7**, **19-21**.
4. K_m_ curve for Boc-QAR-AMC using full-length TMPRSS2.
5. IC_50_ inhibition curves of full-length TMPRSS2 (**Table 2**).
6. Cell-based enzyme activity in HEK-293 cells. Acute toxicity of MM3122 (**4**) data. Activity of **1** and **2** and Camostat using Vero cells in pseudotype and chimeric VSV-SARS-CoV-2 viruses.

