## Supplementary material for "A novel class of TMPRSS2 inhibitors potently block SARS-CoV-2 and MERS-CoV viral entry and protect human epithelial lung cells": Supplemetary Material

**Table of Contents**

1. Single point inhibition data using recombinant protease domain/FRET substrate for **4-6**.
2. Protease selectivity data for **1** and **2**.
3. Synthesis and NMR and HPLC-MS spectra of new compounds **2**, **4-7**, **19-21**.
4. $K_m$ curve for Boc-QAR-AMC using full-length TMPRSS2 (3 nM).
5. $IC_{50}$ inhibition curves of full-length TMPRSS2/Boc-QAR-AMC (**Table 2**).
6. Cell-based enzyme activity in HEK-293 cells. Acute toxicity of MM3122 (**4**) data. Activity of **1** and **2** and Camostat using Vero cells in pseudotype and chimeric VSV-SARS-CoV-2 viruses.

**1. Single point inhibition data using recombinant protease domain and FRET peptide.**

Bar Graph Legend: Compounds (300 µM) were tested against the protease domain of Human Recombinant TMPRSS2 (3 nM) having (MCA)K-KARSAFA-K(Dnp) as substrate (1nM)

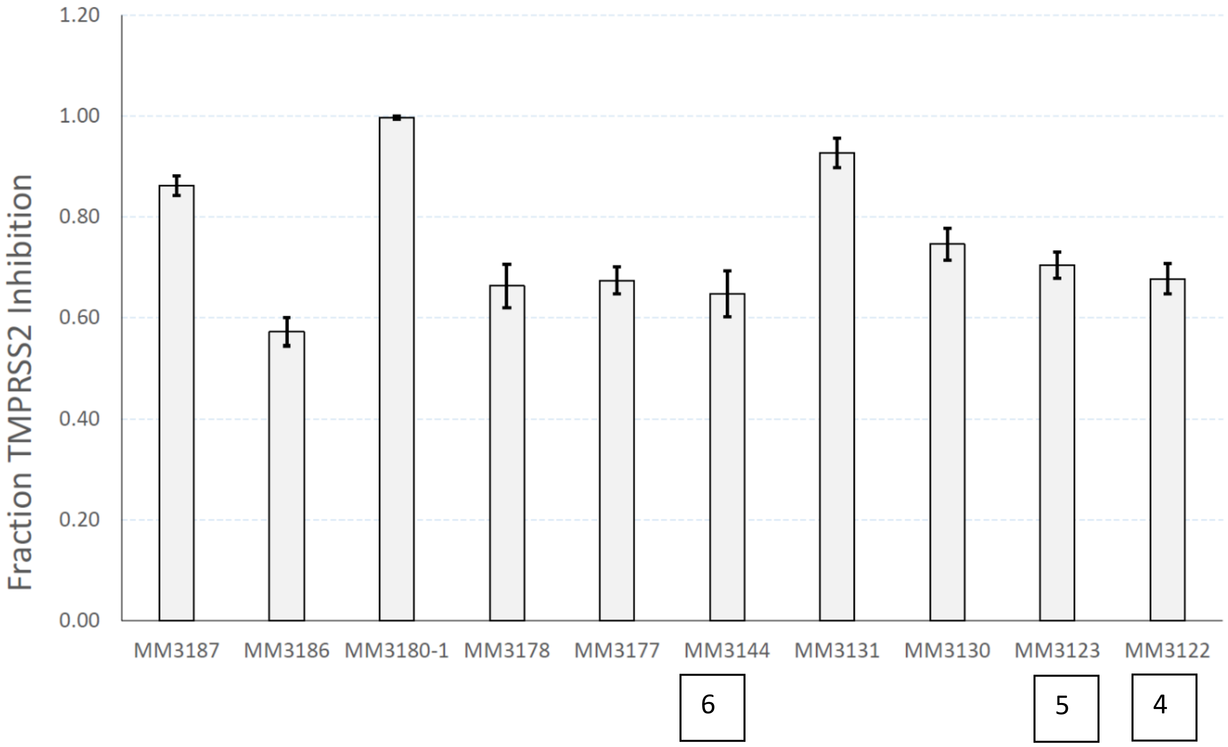

#### 2. Enzyme selectivity data of ZFH7116 (1), VD2173 (2), Camostat and Nafamostat from a panel 43 serine and cysteine proteases. [4]

**Table 1. Protease selectivity of ZFH7116 (1) and VD2173 (2).** NA = not active; all data provided in supplementary material.

| Target | VD2173 IC50 (M) | ZFH7116 IC50 (M) | Camostat IC50 (M) | Nafamostat IC50 (M) |
|---|---|---|---|---|
| Cathepsin B | 2.24E-07 | NA | NA | NA |
| Cathepsin H | 1.04E-05 | NA | NA | NA |
| Cathepsin L | NA | 7.14E-06 | NA | NA |
| Cathepsin S | 1.99E-06 | **8.95E-08** | >1.00E-05 | NA |
| Cathepsin V | NA | 1.00E-05 | NA | NA |
| Chymotrypsin | NA | >2.00E-05 | NA | NA |
| FVIIa | 6.15E-06 | 2.54E-06 | 3.55E-06 | 2.75E-07 |
| FXa | 4.89E-07 | **8.66E-08** | 9.91E-06 | 1.11E-06 |
| FXIa | **7.89E-09** | **1.60E-08** | 3.46E-09 | 8.56E-10 |
| Kallikrein 1 | 1.98E-05 | 2.28E-07 | >1.00E-05 | 2.39E-06 |
| Kallikrein 5 | 3.27E-07 | **8.07E-08** | 1.01E-06 | 6.37E-07 |
| Kallikrein 12 | 5.85E-06 | 5.60E-07 | 1.31E-06 | 3.59E-07 |
| Kallikrein 13 | 1.47E-06 | 5.99E-07 | 8.45E-07 | 3.02E-07 |
| Kallikrein 14 | **4.86E-08** | **1.95E-08** | 9.99E-07 | 2.15E-09 |
| Matriptase 2 | **<1.02E-09** | **<1.02E-09** | 7.80E-09 | <5.08E-10 |
| Papain | **2.67E-08** | 1.97E-05 | NA | NA |
| Plasma Kallikrein | **1.39E-08** | **2.34E-09** | 8.36E-10 | <5.08E-10 |
| Plasmin | 1.73E-06 | 1.34E-07 | 5.62E-09 | 1.04E-09 |
| Proteinase A | 7.59E-06 | 1.88E-06 | NA | NA |
| Proteinase K | **2.21E-08** | **5.19E-09** | NA | NA |
| Thrombin a | 2.14E-07 | 1.33E-06 | 3.62E-06 | 3.03E-07 |
| Trypsin | **<1.02E-09** | **<1.02E-09** | 5.24E-10 | <5.08E-10 |
| Tryptase b2 | **1.10E-09** | **<1.02E-09** | <5.08E-10 | <5.08E-10 |
| Tryptase g1 | **2.34E-09** | **<1.02E-09** | <5.08E-10 | <5.08E-10 |
| Urokinase | **5.01E-09** | 7.82E-06 | 1.64E-08 | <5.08E-10 |

Full protease selectivity data of ZFH7116 (1) and VD2173 (2).

**Report of Protease Profiling for:** Washington Univ. St. Louis **Quotation #** 20200430-WUSM-JJ-Pro-RV02

Two compounds were received as powder stock and resuspended to 10 mM in DMSO.

| Compound ID | Concentration (mM) | Original Stock Volume (uL) | Exact weight (mg) | MW |
|---|---|---|---|---|
| VD2173 | 10 | 382 | 3.0 | 785.84 |
| ZFH7116 | 10 | 349 | 3.2 | 918.00 |

The compound was tested in a 10-dose IC50 with a 3-fold serial dilution starting at 20 uM against 43 proteases.

Control compounds were tested in a 10-dose IC50 with 3-fold serial dilution starting at 10 uM*.
(*Start at different concentrations for some enzymes )

Compound fluorescence : Compounds exhibit no fluorescent background that could interfere with the assay.

The protease activities were monitored as a time-course measurement of the increase in fluorescence signal from fluorescently-labeled peptide substrate, and initial linear portion of slope (signal/min) was analyzed.

Data pages include slope, % Enzyme Activity (No inhibitor control as 100% Activity), curve fit, and IC50.

The obtained IC50 values are summarized in the table below.
(Curve fits were performed when the activities at the highest concentration of compounds were less than 65%.)

Summary Table:

| | | Compound IC50 (M) | | | |
|---|---|---|---|---|---|
| | Target: | VD2173 | ZFH7116 | Control Compound IC50 (M) | Control compound ID |
| 1 | Calpain 1 | | | 4.71E-08 | E64 |
| 2 | Caspase 1 | | | 2.92E-08 | IETD-CHO |
| 3 | Caspase 2 | | | 5.47E-07 | IETD-CHO |
| 4 | Caspase 3 | | | 1.07E-09 | DEVD-CHO |
| 5 | Caspase 4 | >2.00E-05 | | 2.13E-06 | IETD-CHO |
| 6 | Caspase 5 | | | 1.15E-08 | IETD-CHO |
| 7 | Caspase 6 | | >2.00E-05 | 1.24E-08 | DEVD-CHO |
| 8 | Caspase 7 | | | 2.31E-09 | DEVD-CHO |
| 9 | Caspase 8 | | | 2.83E-09 | IETD-CHO |
| 10 | Caspase 9 | | | 2.64E-08 | IETD-CHO |
| 11 | Caspase 10 | | | 9.17E-09 | IETD-CHO |
| 12 | Caspase 11 | | | 5.67E-07 | IETD-CHO |
| 13 | Caspase 14 | | | 6.82E-08 | WEHD-CHO |
| 14 | Cathepsin B | 2.24E-07 | | 7.33E-09 | E64 |
| 15 | Cathepsin C | | | 1.18E-06 | E64 |
| 16 | Cathepsin G | | | 4.04E-06 | Chymostatin |
| 17 | Cathepsin H | 1.04E-05 | | 3.56E-08 | E64 |
| 18 | Cathepsin L | | 7.14E-06 | 1.24E-08 | E64 |
| 19 | Cathepsin S | 1.99E-06 | 8.95E-08 | 1.73E-09 | E64 |
| 20 | Cathepsin V | | 1.00E-05 | 3.89E-09 | E64 |
| 21 | Chymase | | | 1.01E-08 | Chymostatin |
| 22 | Chymotrypsin | | >2.00E-05 | 1.41E-09 | Chymostatin |
| 23 | Elastase | | | 4.83E-09 | Sivelestat |
| 24 | FVIIa | 6.15E-06 | 2.54E-06 | 5.75E-08 | PCI 27483 |
| 25 | FXa | 4.89E-07 | 8.66E-08 | 1.86E-06 | Gabexate mesylate (GM) |
| 26 | FXIa | 7.89E-09 | 1.60E-08 | 2.39E-07 | Gabexate mesylate (GM) |
| 27 | Kallikrein 1 | 1.98E-05 | 2.28E-07 | 4.61E-06 | Leupeptin |
| 28 | Kallikrein 5 | 3.27E-07 | 8.07E-08 | 4.94E-06 | Gabexate mesylate (GM) |
| 29 | Kallikrein 7 | | | 4.31E-05 | Gabexate mesylate (GM) |
| 30 | Kallikrein 12 | 5.85E-06 | 5.60E-07 | 9.43E-08 | Gabexate mesylate (GM) |
| 31 | Kallikrein 13 | 1.47E-06 | 5.99E-07 | 1.22E-05 | Gabexate mesylate (GM) |
| 32 | Kallikrein 14 | 4.86E-08 | 1.95E-08 | 6.51E-07 | Gabexate mesylate (GM) |
| 33 | Matriptase 2 | <1.02E-09 | <1.02E-09 | 3.91E-07 | Gabexate mesylate (GM) |
| 34 | Papain | 2.67E-08 | 1.97E-05 | 2.40E-10 | E64 |
| 35 | Plasma Kallikrein | 1.39E-08 | 2.34E-09 | 1.48E-07 | Gabexate mesylate (GM) |
| 36 | Plasmin | 1.73E-06 | 1.34E-07 | 2.65E-07 | Gabexate mesylate (GM) |
| 37 | Proteinase A | 7.59E-06 | 1.88E-06 | 2.44E-04 | Leupeptin |
| 38 | Proteinase K | 2.21E-08 | 5.19E-09 | 4.51E-08 | Proteinase K inhibitor |
| 39 | Thrombin a | 2.14E-07 | 1.33E-06 | 1.57E-06 | Gabexate mesylate (GM) |
| 40 | Trypsin | <1.02E-09 | <1.02E-09 | 3.07E-08 | Gabexate mesylate (GM) |
| 41 | Tryptase b2 | 1.10E-09 | <1.02E-09 | 9.70E-09 | Gabexate mesylate (GM) |
| 42 | Tryptase g1 | 2.34E-09 | <1.02E-09 | 1.33E-08 | Gabexate mesylate (GM) |
| 43 | Urokinase | 5.01E-09 | 7.82E-06 | 2.44E-08 | Gabexate mesylate (GM) |

* Empty cells indicate no inhibition or compound activity that could not be fit to an IC50 curve

* IC50 value higher than 2.00E-05 M is estimated based on the best curve

* IC50 value lower than 1.02E-09 M is estimated based on the best curve

Report of Protease Profiling for: Washington Univ. St. Louis Quotation # 20200604-WUSM-JJ-PRO

2 compounds were located at RBC as 10 mM DMSO stock.

| Compound ID | Concentration (mM) | Original Stock Volume (uL) | Exact weight (mg) | MW |
|---|---|---|---|---|
| VD2173 | 10 | 382 | 3.0 | 785.84 |
| ZFH7116 | 10 | 349 | 3.2 | 918.00 |

The compounds were tested in a 10-dose IC50 with a 3-fold serial dilution starting at 20 uM against 1 protease.

Control compounds were tested in a 10-dose IC50 with 3-fold serial dilution starting at 1 uM.
(*Start at different concentrations for some enzymes )

Compound fluorescence : Compounds exhibit no fluorescent background that could interfere with the assay.

The protease activities were monitored as a time-course measurement of the increase in fluorescence signal from fluorescently-labeled peptide substrate, and initial linear portion of slope (signal/min) was analyzed.

Data pages include slope, % Enzyme Activity (No inhibitor control as 100% Activity), curve fit, and IC50.

The obtained IC50 values are summarized in the table below.
(Curve fits were performed when the activities at the highest concentration of compounds were less than 65%.)

Summary Table:

| | | Compound IC50 (M) | | | |
|---|---|---|---|---|---|
| | Target: | VD2173 | ZFH7116 | Control Compound IC50 (M) | Control compound ID |
| 1 | Furin | | | 1.10E-09 | Furin Inhibitor 1 |

* Empty cells indicate no inhibition or compound activity that could not be fit to an IC50 curve

#### 3. Synthesis and NMR and HPLC-MS spectra of new compounds 2, 4-7, 19-21.

##### Synthesis of peptidyl ketobenzothiazoles (kbts), 4-7

We have previously published methodology for the synthesis of the kbt peptide inhibitors [1]. As shown in **Scheme 1**, we first construct the tripeptide (e.g. **6**) or tetrapeptide (e.g. **4**) on the 2-chlorotrityl resin using standard Fmoc-solid phase peptide synthesis protocols including HBTU for amide bond coupling and 20% piperidine for Fmoc deprotection steps. The peptide is then capped with an acetyl group using acetic anhydride. Cleavage of the 2-chlorotrityl resin without removal of the protecting groups is accomplished using HFIP, then H-Arg(Mtr)-kbt is installed with HATU or EDC/HOBt in DMF. Final deprotection of the amino acid sidechains using TFA:water:thioanisole (95:2.5:2.5 %v/v) generates the target compounds which are purified by reverse phase prep HPLC.

**Scheme 1. Synthesis of acyclic P4-P1 tetrapeptide ketobenzothiazoles (kbts), 4-7.**

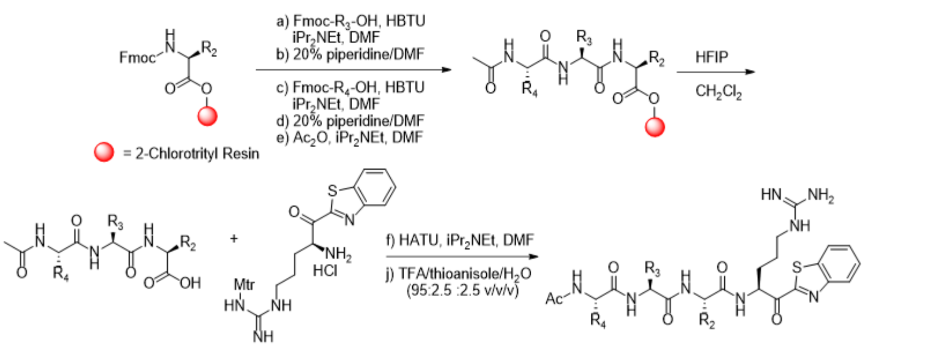


**Scheme 2. Cycloamide synthesis (2, 19 and 20)**

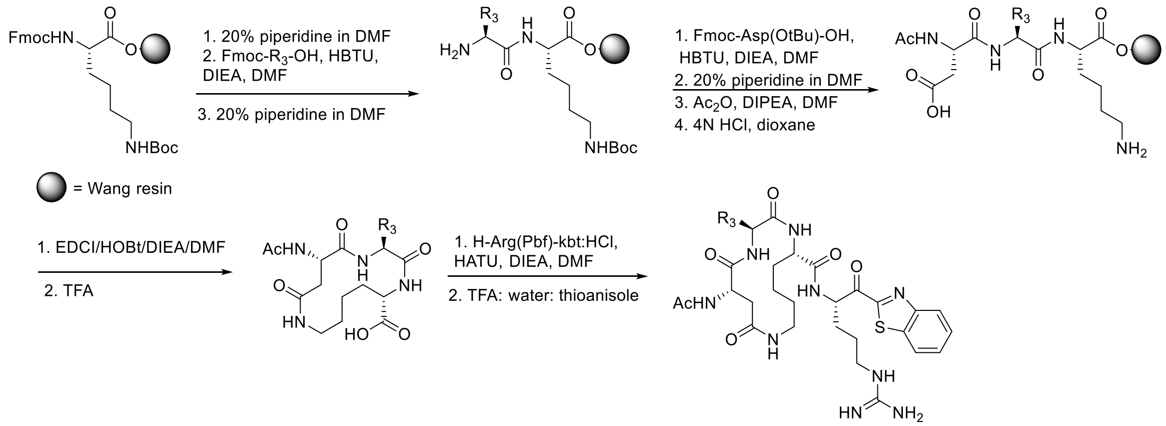


Cyclic peptide VD2173 (**2**) was synthesized as described in another publication[2] but follows the generic procedure outlined in **Scheme 2** in which is also used for the construction of **19** and **20**. Synthesis of the tripeptide cyclization precursors are synthesized on Wang resin using standard Fmoc solid phase peptide synthesis (SPPS) using 2HBTU for coupling steps and 20% piperidine for Fmoc removal steps. After the final Fmoc deprotection, the tripeptide is acetylated and then the Asp and Lys protecting groups are removed with 4N HCl. Cyclization is performed using EDCI and HOBt on the resin followed by TFA cleavage, then a final HATU coupling with H-Arg-(Pbf)-kbt followed by Arg deprotection to give the cyclic peptides which are purified by reverse phase prep HPLC.

**Scheme 3. Synthesis of cyclic peptide aryl ether 21**

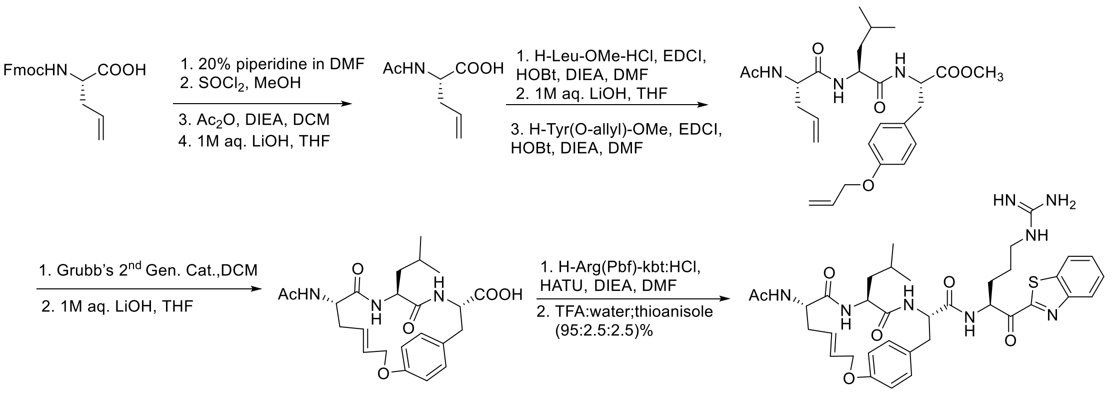


Compound **21** was prepared from Fmoc-allylglycine as shown in **Scheme 3**. N-Deprotection followed by esterification and acetylation gives Ac-allylglycine which is then coupled to H-Leu-OMe using EDC/HOBt. The resulting dipeptide ester is hydrolyzed with LiOH and then coupled to H-Tyr(Oallyl)-OMe once again with EDC/HOBt to yield the cyclization precursor. Olefin metathesis cyclization is accomplished with Grubbs 2$^{nd}$ generation catalyst to give key aryl allyl ether cyclic peptide intermediate. Ester hydrolysis followed by amide coupling to H-Arg(Pbf)-kbt and final deprotection with TFA as before gives **21** which is purified by prep HPLC.

**General synthesis, purification, and analytical chemistry procedures.** Starting materials, reagents, and solvents were purchased from commercial vendors unless otherwise noted. $^1$H NMR spectra were measured on a Varian 400 MHz NMR instrument. The chemical shifts were reported as δ ppm relative to TMS using residual solvent peak as the reference unless otherwise noted. The following abbreviations were used to express the multiplicities: s = singlet; d = doublet; t = triplet; q = quartet; m = multiplet; br = broad. High-performed liquid chromatography (HPLC) was carried out on GILSON GX-281 using Waters C18 5µM, 4.6*50mm and Waters Prep C18 5µM, 19*150mm reverse phase columns, eluted with a gradient system of 5:95 to 95:5 acetonitrile:water with a buffer consisting of 0.05% TFA. Mass spectra (MS) were performed on HPLC/MSD using electrospray ionization (ESI) for detection. All reactions were monitored by thin layer chromatography (TLC) carried out on Merck silica gel plates (0.25 mm thick, 60F254), visualized by using UV (254 nm) or dyes such as $KMnO_4$, *p*-Anisaldehyde and CAMA (Cerium Ammonium Molybdate or Hanessian's Stain). Silica gel chromatography was carried out on a Teledyne ISCO CombiFlash purification system using pre-packed silica gel columns (12g to 330g sizes). All compounds used for biological assays are greater than 95% purity based on NMR and HPLC by absorbance at 220 nm and 254 nm wavelengths.

**General procedure for synthesis of acyclic peptides**[1]

**Solid phase peptide coupling and deprotection:** Into a reaction vial (with a fritted glass filter) under nitrogen containing H-Leu-2-Cl trityl or H-Phe-2-Cl trityl resin (0.714 g, 0.5 mmol) was added DMF/$CH_2Cl_2$ (15/15 mL). The mixture was shaken at RT for 30 min and then filtered. The resin was washed with DMF (10 mL) 2 times. A mixture of Fmoc-AA-OH (2.5 mmol) in DMF (20 mL), HBTU (0.853 g, 2.25 mmol) and $^{i}Pr_2NEt$ (0.87 mL, 5 mmol) was stirred at RT for 10 min and then added to the resin. The resultant heterogeneous mixture was shaken at RT overnight and then filtered. The resin was washed with DMF (20 mL x 4), dried and then piperidine/DMF (20% v/v, 30 mL) was added. The mixture was shaken for 1-4 h at RT, then filtered and was washed with DMF (10 mL x 4). Following the Fmoc deprotection of the dipeptide, the dipeptide is carried on to the next step or coupling of another Fmoc-AA-OH is performed in an identical fashion as described above and then subsequently a final Fmoc deprotection to the tripeptide.

**Acetyl capping and cleavage from resin:** The peptide-containing resin was suspended in 30 mL of 0.5 M $Ac_2O$ and 1 M $^{i}Pr_2NEt$ in DMF and shaken at RT for 1 h. The reaction was filtered, and resin washed with DMF (10 mL x 4) followed by $CH_2Cl_2$ (10 mL x 4). The resin was then suspended in 30 ml of 25% v/v HFIP/ $CH_2Cl_2$ and shaken for 1 h. The reaction was filtered, and the filtrate was concentrated and dried *in vacuo*.

**Coupling of Arg (Pbf)-kbt:HCl and final deprotection.** To crude peptide acid (400 mg, 1.0 mmol) dissolved in dry DMF (10 mL) under a nitrogen atmosphere at 0 °C was added HATU (456 mg, 1.20 mmol) followed by stirring for 15 min. Next, Arg(Pbf)-kbt:HCl (638mg; 1.10 mmol) and *i*$Pr_2NEt$ (0.87 mL, 5.0 mmol) were added to the reaction at 0 °C. The reaction was allowed to reach room temperature and then stirred for an additional 2-3 h. DMF was removed under vacuum and water (250 mL) was added to the residue. The precipitate formed was filtered and washed with water (2 x 50 mL) then dried under vacuum. The precipitate was suspended in 10 mL TFA/thioanisole/water (95:2.5:2.5 *v/v/v*) and stirred for2 h at RT. The solvent was removed, and cold ether (100 mL) was added. The resulting precipitate was collected by centrifugation and the crude product was purified by HPLC ($C_{18}$, 15 x 150 mm column; eluent: acetonitrile/water (0.05% TFA) to give the final compound.

**Ac-WFR-kbt, 8.** $^{1}H$ NMR (399 MHz, DMSO-*d*$_6$) d ppm 1.48 (br. s., 1 H), 1.54 - 1.69 (m, 1 H), 1.74 (m, 4 H), 2.70 - 2.89 (m, 2 H), 2.90 - 3.22 (m, 4 H), 4.37 - 4.74 (m, 1 H), 5.41 - 5.58 (m, 1 H), 6.86 - 7.10 (m, 4 H), 7.11 - 7.36 (m, 8 H), 7.40 - 7.57 (m, 2 H), 7.62 - 7.76 (m, 2 H), 7.93 - 8.12 (m, 2 H), 8.22 - 8.34 (m, 2 H), 8.60 (t, *J*=7.59 Hz, 1 H), 10.76 (s, 1 H). ESI-MS [M+H]+ calcd for for $C_{35}H_{38}N_8O_4S^+$ 666.80, found 667.5.

**Ac-dWFR-kbt, 13.** $^1$H NMR (399 MHz, DMSO-$d_6$) d ppm 1.56 - 2.07 (m, 6 H), 1.93 - 2.04 (m, 1 H), 2.52 - 2.81 (m, 4 H), 2.96 - 3.08 (m, 1 H), 4.39 - 4.50 (m, 1 H), 4.57 - 4.70 (m, 1 H), 5.46 - 5.58 (m, 1 H), 6.90 - 7.09 (m, 4 H), 7.09 - 7.36 (m, 8 H), 7.44 - 7.59 (m, 2 H), 7.62 - 7.76 (m, 2 H), 7.99 (d, *J*=7.79 Hz, 1 H), 8.28 (t, *J*=8.37 Hz, 2 H), 8.47 (d, *J*=8.95 Hz, 1 H), 8.63 (d, *J*=6.62 Hz, 1 H), 10.74 (s, 1 H). ESI-MS [M+H]+ calcd for $C_{35}H_{38}N_8O_4S^+$ 666.80, found 667.5.

**Ac-QFR-kbt, 6.** $^1$H NMR (400 MHz, DMSO-$d_6$) d ppm 0.85 - 1.23 (m, 10 H), 1.23 - 1.49 (m, 4 H), 2.15 (dd, *J*=13.89, 8.41 Hz, 1 H), 2.30 (dd, *J*=13.69, 6.26 Hz, 1 H), 2.37 - 2.48 (m, 2 H), 3.41 - 3.52 (m, 1 H), 3.79 - 3.89 (m, 1 H), 4.86 - 4.96 (m, 1 H), 6.24 - 6.42 (m, 7 H), 6.84 (quin, *J*=7.53 Hz, 3 H), 7.33 (d, *J*=7.43 Hz, 1 H), 7.37 - 7.45 (m, 3 H), 7.73 (d, *J*=7.04 Hz, 1 H). ESI-MS [M+H]+ calcd for $C_{29}H_{36}N_8O_5S^+$ 608.72, found 609.5.

**Ac-IQFR-kbt, 7.** H NMR (400 MHz, DMSO-$d_6$) d ppm 0.70 - 0.84 (m, 7 H) 1.00 - 1.14 (m, 1 H) 1.31 - 1.48 (m, 1 H) 1.54 - 1.84 (m, 7 H) 1.84 - 2.13 (m, 7 H) 2.72 - 2.84 (m, 1 H) 2.95 - 3.07 (m, 1 H) 3.10 - 3.21 (m, 2 H) 4.08 (t, *J*=6.85 Hz, 1 H) 4.12 - 4.21 (m, 1 H) 4.51 - 4.64 (m, 1 H) 5.45 - 5.55 (m, 1 H) 6.81 (br. s., 1 H)7.10 - 7.31 (m, 6 H) 7.49 - 7.57 (m, 1 H) 7.64 - 7.74 (m, 2 H) 7.95 (t, *J*=9.00 Hz, 2 H) 8.15 (d, *J*=7.83 Hz, 1 H) 8.28 (t, *J*=8.61 Hz, 2 H) 8.56 (d, *J*=5.87 Hz, 1 H). ESI-MS [M+H]+ calcd for $C_{35}H_{47}N_9O_6S^+$ 721.88, found 722.6.

**Ac-GQFR-kbt, 4 (MM122).** $^1$H NMR (400 MHz, DMSO-*d6*) δ ppm 1.51 - 1.69 (m, 3 H), 1.70 - 1.83(m, 2 H), 1.87 (s, 3 H), 1.91 - 2.11 (m, 3 H), 2.71 - 2.86 (m, 1 H), 2.98 - 3.08 (m, 1 H), 3.11 - 3.22 (m, 2 H), 3.68 (d, *J*=5.48 Hz, 2 H), 4.10 - 4.23 (m, 1 H), 4.55 (br. s., 1 H), 5.50 (br. s., 1 H), 6.80 (br. s., 1 H), 7.09 - 7.32 (m, 7 H), 7.53 (br. s., 1 H), 7.63 - 7.75 (m, 2 H), 8.02 (d, *J*=8.22 Hz, 1 H), 8.09 - 8.21 (m, 2 H), 8.23 - 8.34 (m, 2 H), 8.52 (d, *J*=6.26 Hz, 1 H). ). ESI-MS [M+H]+ calcd for $C_{31}H_{39}N_9O_6S^+$ , 665.77, found 666.50.

**Ac-PQFR-kbt, 5.** $^1$H NMR (400 MHz, DMSO-$d_6$) δ ppm 0.87 - 1.54 (m, 19 H), 2.09 -2.24 (m, 1 H), 2.37 - 2.49 (m, 3 H), 2.76 - 2.89 (m, 1 H), 2.96 - 3.09(m, 1 H), 3.35 - 3.57 (m, 2 H), 3.78 - 3.94 (m, 1 H), 4.90 (br. s., 1 H), 6.25 - 6.53 (m, 6 H), 6.76 - 6.90 (m, 2 H), 7.23 (d, J=7.43 Hz, 1 H), 7.33 (d, J=7.83 Hz, 1 H), 7.37 - 7.48 (m, 2 H), 8.44 (d, J=5.48 Hz, 1H). ). ESI-MS [M+H]+ calcd for $C_{34}H_{43}N_9O_6S^+$ 705.84, found 706.50.

**H-dWFR-kbt, 14**:

**Boc-dWF-OMe**: Boc-D-Trp-OH (1 g, 3.28 mmol) and HCl.Phe-OMe (0.708g, 3.28 mmol) was taken in dry dichloromethane (10 mL) under nitrogen atmosphere and the reaction mixture was cooled to 0 $^oC$ and N,N-diisopropylethylamine (1.7 mL, 9.84 mmol) and propylphosphonic anhydride (1.9 mL, 3.28 mmol, 50% solution in EtOAc) were added to the solution drop wise respectively. The reaction mixture was then stirred at 25 $^oC$ under nitrogen atmosphere for 1 hour and the completion of the reaction was confirmed by LC-MS monitoring. On completion the reaction mixture was diluted with 10 mL dichloromethane and washed with 10% citric acid solution, saturated sodium bicarbonate solution and brine respectively. The organic layer was dried over sodium sulfate and concentrated under reduced pressure. The crude product was triturated with hexane to obtain the title product in pure form as a white solid. Yield: 1.3 g (92.8%). Chemical formula: $C_{23}H_{33}N_3O_5$, Exact Mass: 465.55, MS(ESI): found: $[M + Na]^+$, 488.58.

**Boc-dWF-OH**: Boc-WF-OMe (0.130 g, 0.279 mmol) was taken in a 1:1 mixture of THF and water and LiOH.$H_2O$ (0.035g, 0.873 mmol) was added to it. The reaction mixture was stirred for 30 minutes at 25 $^oC$ and the completion of the reaction was confirmed by LCMS monitoring. On completion, the THF was evaporated under reduced pressure and the remaining water layer was cooled to 0 $^oC$. The water layer was then brought to pH 6.5 by slow addition of 0.5 M HCl solution in water. The crude product precipitates out on addition of HCl and it was isolated by filtration. The crude product was dried under reduced pressure and triturated with diethyl ether to obtain the pure title product in pure form as white solid. Yield: 0.102g (80%). Exact Mass: 451.21, MS(ESI): found: $[M + H]^+$, 452.26.

**Boc-dWFR(Mtr)-kbt**: Boc-dWF-OH (35 mg, .077 mmol) and HATU (43.9 mg, 0.115 mmol) was taken in dry DMF under nitrogen atmosphere and the reaction mixture was cooled to 0 $^oC$. N,N-diisopropylethylamine (0.04mL, 0.231 mmol) was then added drop wise to the reaction mixture and the reaction mixture was allowed to stir for 15 minutes followed by addition of HCl.Arg(Mtr)-kbt (41.75 mg, .077 mmol). The reaction mixture was stirred for 12 hours at 25 $^oC$ under nitrogen atmosphere and the completion of the reaction was confirmed by LC-MS monitoring. On completion, the reaction mixture was diluted with EtOAc and washed with 10% citric acid solution, saturated sodium bicarbonate solution and brine respectively. The organic layer was dried over sodium sulfate and concentrated under reduced pressure. The crude product was directly taken to the next step without further purification. Chemical formula: $C_{48}H_{56}N_8O_8S_2$, Exact Mass: 936.37, MS(ESI): found: $[M + H]^+$, 937.17.

**H-dWFR-kbt (14)**: Boc-dWFR(Mtr)-kbt (85 mg, crude product from previous step) was taken in 5 mL TFA:thioanisole:$H_2O$ (95:2.5:2.5) and the reaction mixture was stirred for 6 hours at 25 $^oC$. The completion of the reaction was confirmed by LC-MS monitoring. On completion, the reaction mixture was concentrated under reduced pressure and triturated with diethyl ether to obtain the crude product as brown solid. The crude product was then subjected to reverse phase semi-preparative HPLC (Stationary phase: C18 column, mobile phase: $H_2O$-Acetonitrile with 0.1% TFA in each, 15-65% Acetonitrile in $H_2O$ gradient for 20 minutes) to obtain the pure title product Yield: 25 mg (42% over two steps). Chemical formula: $C_{33}H_{36}N_8O_3S$, $^1H$ NMR (400 MHz, METHANOL-d4) δ ppm 8.21 (d, *J* = 6.65 Hz, 1H), 8.11 (d, *J* = 7.43 Hz, 1H), 7.57 - 7.67 (m, 4H), 7.12 - 7.36 (m, 10H), 7.01 - 7.08(m, 2H), 6.84 (s, 1H), 3.48 (d, *J* = 1.96 Hz, 1H), 3.13 (d, *J* = 6.65 Hz, 4H), 3.01 - 3.07 (m, 1H), 2.75 - 2.84 (m, 1H), 2.65 (s, 2H), 2.03 (s, 5H), 1.41(d, *J* = 8.61 Hz, 1H)Exact Mass: 624.26, MS (ESI): found: $[M + H]^+$, 625.5.

**H-dWFR-kbt-COOH, 15:** The title compound was synthesized using the same procedure as **14** using H-Arg(Mtr)-kbt-COOH:HCl[1]. Yield: 24 mg (43% over two steps). Chemical formula: $C_{34}H_{36}N_8O_5S$, $^1H$ NMR (400 MHz,

METHANOL-d4) δ ppm 10.58 (br. s., 0H), 8.82 (s, 1H), 8.22 - 8.32 (m, 2H), 7.70 (dd, *J* = 8.02, 16.24 Hz, 0H), 7.57 - 7.64 (m, 1H),7.37 (d, *J* = 6.26 Hz, 1H), 7.22 (d, *J* = 13.30 Hz, 0H), 7.11 - 7.18 (m, 2H), 7.02 - 7.10 (m, 1H), 5.55 - 5.67 (m, 1H), 4.47 - 4.58 (m, 0H), 4.35 (dd, *J*= 5.09, 10.17 Hz, 0H), 4.05 - 4.31 (m, 2H), 3.37 - 3.55 (m, 1H), 3.07 - 3.27 (m, 3H), 2.65 (s, 1H), 2.18 (dd, *J* = 6.06, 13.11 Hz,1H), 1.57 - 1.95 (m,3H), 1.16 - 1.52 (m, 2H), 1.06 (td, *J* = 7.14, 13.89 Hz, 1H), 0.91 - 1.00 (m, 2H), 0.65 - 0.81 (m, 6H) Exact Mass: 668.253, MS (ESI): found: $[M + H]^+$, 669.5.

**Ac-Cyclo(DLK)-R- ketobenzothiazole (2, VD2173).** Into a reaction vessel (with fritted glass for resin support) containing Fmoc-L-Lys(Boc) Wang resin (5 g, 1.7 mmol), DCM (40 mL) was added. The mixture was shaken at RT for 15 min and then filtered. To the dry resin was added piperidine/DMF (20% *v/v*, 40 mL) and the mixture was shaken for 30 min at RT, then filtered. The resin was washed with DMF (2 x 30mL) and DCM (2 x 30 mL). Fmoc-Leu-OH (1.8 g, 5.1 mmol), HBTU (2.25 g, 5.95 mmol), *i*$Pr_2NEt$ (1.31 g, 10.2 mmol), and DMF (50 mL) were added to the vessel and shaken for 12 h, then filtered. The resin was washed with DCM (2 x 40 mL) and DMF (2 x 40 mL), then piperidine/DMF (20% *v/v*, 40 mL) was added and the reaction was shaken for 30 min at RT, then filtered. The resin washed with DCM (2 x 30 mL) and DMF (2 x 30mL). Fmoc-Asp(OtBu)-OH (2.10 g, 5.1 mmol), HBTU (2.25 g, 5.95 mmol), *i*$Pr_2NEt$ (1.31 g, 10.2 mmol), and DMF (50 mL) were added to the vessel and shaken for 12 h, then filtered. The resin was washed with DCM (2 x 40 mL) and DMF (2 x 40 mL). The peptide resin was then suspended in a solution of $Ac_2O$ (1.04 g, 10.2 mmol), and *i*$Pr_2NEt$ (3.07 g, 23.8 mmol) in 40 mL of DMF. The mixture was shaken at RT for 1-2 h, filtered and resin washed with DCM (2 x 40 mL) followed by DMF (2 x 40 mL). To the resin was added 40 mL of dry 4M HCl in 1, 4-dioxane followed by shaking for 30-40 min. at RT. The reaction was filtered, and the resin washed with DCM (2 x 40 mL) followed by DMF (2 x 40 mL). EDCI (0.98 g, 5.1 mmol), HOBt (0.78 g, 5.1 mmol), *i*$Pr_2NEt$ (1.1 g, 8.5 mmol), and DMF (80 mL) were added to the resin and the resulting mixture was shaken for overnight at RT. The mixture was filtered and the resin and washed with DCM (2 x 40 mL) followed by DMF (2 x 40 mL). To the acetyl capped peptide resin was added TFA (2 x 35 mL) and shaken for 30 min. The mixture was filtered, and the resin washed with DCM (2 x 40 mL). The filtrate was concentrated, and cold ether was added to the residue yielding the crude product as a precipitate which was purified by flash chromatography to give an off-white sold (400 mg).

The macrocyclic tripeptide acid (400 mg, 1.0 mmol) was dissolved in dry DMF (10 mL) under a nitrogen atmosphere at 0 °C and HATU (456 mg, 1.20 mmol) was added followed by stirring for 15 min, and then the addition of Arg(Pbf)-kbt:HCl ( 638mg;  1.10 mmol) and *i*$Pr_2NEt$ (0.87 mL, 5.0 mmol) at 0 °C. The reaction is allowed to reach RT and then stirred for 2-3 h. The DMF was removed in vacuo and water (250 mL) was added to the resulting residue. The precipitate formed was filtered and washed with water (2 x 50 mL) and dried. To this precipitate was added 10 mL of TFA/thioanisole/water (95:2.5:2.5 *v/v/v*) and the mixture was stirred for 2 h at RT. The solvent was removed, and then cold ether (100 mL) was added. The crude product was collected by centrifugation. The crude product was purified by HPLC ($C_{18}$, 15 x 150 mm column; eluent: acetonitrile/water (0.05% TFA) to give the title compound as a white solid. Overall yield (20%).1H NMR (400MHz ,DMSO-$d_6$) δ ppm = 8.51 (d, *J* = 6.7 Hz, 1 H), 8.26 (dd, *J* = 8.0, 15.1 Hz, 1 H), 7.98 (d, *J* = 7.4 Hz, 1 H), 7.93 - 7.83 (m, 2 H), 7.73 - 7.63 (m, 2 H), 7.53 (br. s., 1 H), 5.44 - 5.33 (m, 1 H), 4.60 - 4.48 (m, 1 H), 4.29 - 4.18 (m, 1 H), 3.42 (br. s., 4 H), 3.19 - 3.06 (m, 3 H), 2.96 (br. s., 1 H), 1.84 (s, 3 H), 1.78 - 1.69 (m, 1 H), 1.65 - 1.33 (m, 8 H), 1.23 - 1.07 (m, 2 H), 0.89 - 0.74 (m, 7 H). ESI-MS [M+H]+ calcd for $C_{31}H_{46}N_9O_6S^+$ 672.33, found 672.5.

**Ac-Cyclo(DQK)-R- ketobenzothiazole, 20.** Synthesized like VD2173. Overall yield (30%).$^1$H NMR (400MHz, DMSO-$d_6$) δ ppm = 9.19 (s, 1 H), 8.98 - 8.81 (m, 2 H), 8.78 - 8.67 (m, 1 H), 8.60 - 8.50 (m, 1 H), 8.38 - 8.24 (m, 1 H), 8.24 - 8.13 (m, 1 H), 7.89 - 7.81 (m, 1 H), 7.41 - 7.32 (m, 1 H), 7.29 - 7.11 (m, 1 H), 6.21 - 5.98 (m, 2 H), 5.25 - 5.08 (m, 1 H), 4.94 - 4.82 (m, 1 H), 4.05 (br. s., 2 H), 3.79 (t, *J* = 6.1 Hz, 6 H), 3.15 – 3.04 (m, 6 H), 2.69 - 2.56 (m, 3 H), 2.44 - 2.32 (m, 3 H), 2.04 (s, 3 H), 2.25 (br. s., 5 H). ESI-MS [M+H]+ calcd for $C_{30}H_{43}N_{10}O_7S^+$ 687.30, found 687.50.

**Ac-Cyclo(DMK)-R- ketobenzothiazole, 19.** Synthesized like VD2173. Overall yield (27%). $^1$H NMR (400MHz, DMSO-d6) δ ppm = 8.54 (d, *J* = 6.3 Hz, 1 H), 8.31 - 8.07 (m, 2 H), 7.93 (d, *J* = 8.2 Hz, 1 H), 7.68 (br. s., 1 H), 7.49 (br. s., 1 H), 5.40 (br. s., 1 H), 4.51 (br. s., 1 H), 4.22 (br. s., 1 H), 3.15 (br. s., 6 H), 2.00 (d, *J* = 1.2 Hz, 4 H), 1.88 - 1.77 (s, 3 H), 1.58 (br. s., 3 H), 1.39 - 1.03 (m, 16 H). ESI-MS [M+H]+ calcd for $C_{30}H_{43}N_9O_6S_2^+$ 690.28, found 690.40.

**Ac-Cyclo(Allyl-Y)-R- ketobenzothiazole, 21.** Fmoc-(L)-glycine (3.5 g, 10 mmol) stirred in 20% piperidine in DMF (20 mL) for 1 hr. Solvent was removed under reduced pressure, product triturated with DCM and hexanes (1:3), filtered the product and washed with hexanes, dried and used in the next reaction. Above material was dissolved in methanol (10 mL) and cooled the reaction to 0 °C followed by added thionyl chloride (2 mL) dropwise and stirred for 10 min and ice bath was replaced by a water bath, and the reaction mixture heated to ~50 °C for 3 hr while stirring. Removal of the solvent left a white residue which was washed with diethyl ether (100 mL) and collected by vacuum filtration to yield the amino acid methyl ester hydrochloride as a solid (1.7 g). Above ester (500 mg; 3.02 mmol) was taken in DCM (10 mL) and added DIEA (1.58 mL; 9.06 mmol) and $Ac_2O$ (0.86 mL; 9.06 mmol) at RT and stirred for 3 hrs. Solvent was removed under reduced pressure and crude was purified by flash chromatography using EtOAc and Hexanes (1:9). A solution of ester (395 mg; 2.5 mmol) in THF (3 mL) was treated with 1M aqueous LiOH (3 mL) and the reaction mixture was stirred for 3 h at RT, and the absence of starting material was monitored by TLC. After the solvent was evaporated off, the residue was diluted with water and the pH was adjusted to ~3.0 using 5% aq. HCl. The product was extracted with ethyl acetate (2 x 50 mL) and the combined organic layer washe with brine (20 mL), dried over anhydrous $Na_2SO_4$, filtered off and concentrated, which is used in the next step without further purification. N-acetyl allyl glycine acid (167 mg;

1.06 mmol) in DMF (5 mL) was stirred with peptide coupling reagent EDCI/HOBt or HATU (1.3 eq) for 30 min. The reaction was cooled to 0-5 °C and charged with amino acid methyl ester hydrochloride (1.1 eq.) followed by diisopropylethylamine (3.0 eq.). After 15 min, allowed the reaction was brought to RT and stirred overnight. Solvent was removed under reduced pressure and the residue partitioned between EtOAc and 5% aq. HCl. The separated organic layer was washed with aq. 5% HCl, saturated $NaHCO_3$ solution (2x) and brine (1x) then dried over anhydrous $Na_2SO_4$. The crude product was purified by silica gel column chromatography using EtOAc and Hexanes (2:8). A solution of the ester (343 mg; 1.2 mmol) in THF (4 mL) was treated with 1M aqueous LiOH (4 mL). The reaction mixture was stirred for 3 h at RT, and the absence of starting material was monitored by TLC. After the solvent was evaporated off, the residue was diluted with water and the pH was adjusted to ~3.0 using 5% aq. HCl. The crude product was extracted into ethyl acetate (3 x 100 mL). The combined organic layers were washed with brine (25 mL) and dried over anhydrous $Na_2SO_4$. N-acetyl dipeptide acid (135 mg; 0.5 mmol) was stirred with EDCI (1.3 eq) and HOBt (1.3 eq) in DMF (3 mL) for 30 min. The reaction was cooled to 0–5 °C and H-L- O-allyl Tyr-OMe. HCl (130 mg: 0.55 mmol) followed by DIEA (3.0 eq). After 15 min, allowed the reaction to RT and stirred overnight. Solvent was removed under reduced pressure and the residue partitioned between EtOAc and 5% aq. HCl. The separated organic layer was washed with aq. 5% HCl, saturated $NaHCO_3$ solution (2x) and brine (1x) then dried over anhydrous $Na_2SO_4$. The crude product was purified by silica gel column chromatography using EtOAc and Hexanes (3:7).

A solution of acyclic diene precursor (150 mg, 0.3076 mmol) in DCM (280 mL, 0.2 Mol.) degassed for 30 min by purging nitrogen gas and then Grubbs 2$^{nd}$ generation catalyst (26 mg, 10 mol %) was added. The reaction was refluxed for 30 min and then additional Grubbs 2$^{nd}$ generation (13 mg, 5 mol %) was added. The reaction was refluxed for 18 h under nitrogen atmosphere. After depletion of the starting material as monitored by TLC and LCMS, the reaction was cooled to RT and quenched by adding activated charcoal (100 mg) followed by stirring for 1 h. The mixture was filtered through celite bed and washed generously with DCM. The filtrate was concentrated in vacuo and the crude product was purified by silica gel chromatography to yield an off-white solid. A solution of the macrocyclic ester (50 mg: 0.10 mmol) in MeOH (2 mL) was treated with 1M aq. LiOH (2 mL) at RT for 3 hrs, and the absence of starting material was monitored by TLC. After the solvent was evaporated off, the residue was diluted with water and the pH was adjusted to ~3.0 using 5% aq. HCl. The product was extracted with ethyl acetate (3 x 50 mL) and the combined organic layer washe with brine (20 mL), dried over anhydrous $Na_2SO_4$, filtered off and concentrated, which is used in the next step without further purification.

The macrocyclic acid (45 mg, 0.101 mmol) was dissolved in dry DMF (3 mL) under a nitrogen atmosphere at 0 °C and HATU (50 mg, 1.30 mmol) was added followed by stirring for 15 min, and then the addition of Arg(Pbf)-kbt:HCl ( 65 mg;  0.111 mmol) and *i*$Pr_2NEt$ (70 uL, 0.404 mmol) at 0 °C. The reaction is allowed to reach RT and then stirred for 2-3 h. The DMF was removed in vacuo and water (250 mL) was added to the resulting residue. The precipitate formed was filtered and washed with water (2 x 50 mL) and dried. To this precipitate was added 2.5 mL of TFA/thioanisole/water (95:2.5:2.5 *v/v/v*) and the mixture was stirred for 2 h at RT. The solvent was removed, and then cold ether (35 mL) was added. The crude product was collected by centrifugation. The crude product was purified by HPLC ($C_{18}$, 15 x 150 mm column; eluent: acetonitrile/water (0.05% TFA) to give the title compound as a white solid. Overall yield (32%). $^1H$ NMR (400MHz, DMSO-d6) δ ppm = 8.79 (d, *J* = 7.4 Hz, 1 H), 8.33 - 8.25 (m, 2 H), 8.20 (d, *J* = 9.0 Hz, 1 H), 8.13 (d, *J* = 7.4 Hz, 1 H), 7.75 - 7.64 (m, 2 H), 7.53 (br. s., 1 H), 7.07 (d, *J* = 7.8 Hz, 2 H), 6.67 (d, *J* = 7.8 Hz, 1 H), 5.60 - 5.36 (m, 2 H), 4.72 - 4.54 (m, 2 H), 4.37 - 4.25 (m, 1 H), 4.17 (br. s., 1 H), 3.16 (d, *J* = 6.3 Hz, 2 H), 2.98 (d, *J* = 11.7 Hz, 3 H), 1.79 (s, 3 H), 1.64 (br. s., 4 H), 1.41 - 1.32 (m, 3 H), 1.24 - 1.14 (m, 4 H), 0.83 - 0.72 (m, 10 H).

The synthesis of **9**, **10**, **11**, and **18** have been previously reported.[3]

The synthesis of **1**, **3**, **12**, **16**, and **17** are as previously described.[1]

NMR and HPLC-MS spectra of new compounds **4-7**, **2**, **19-21**.

VD2173 (Compound 2)

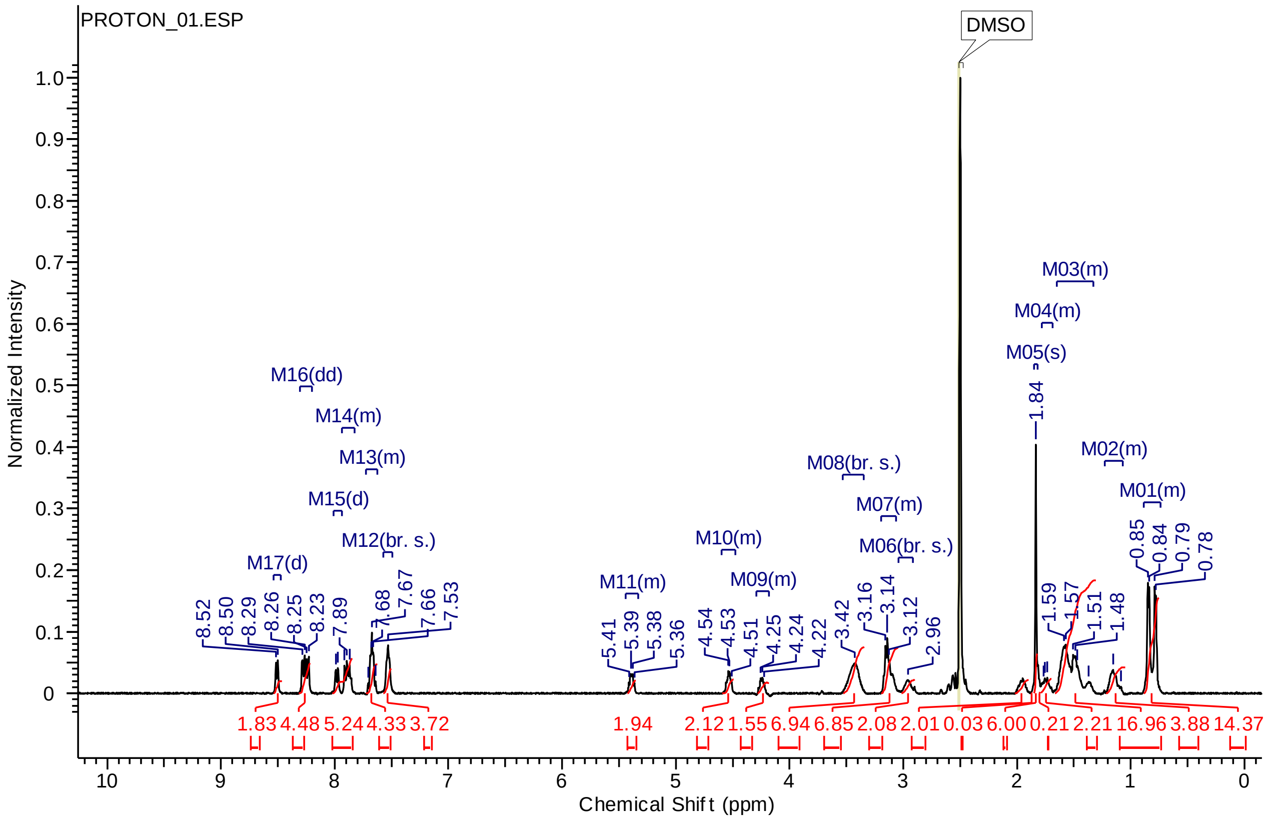


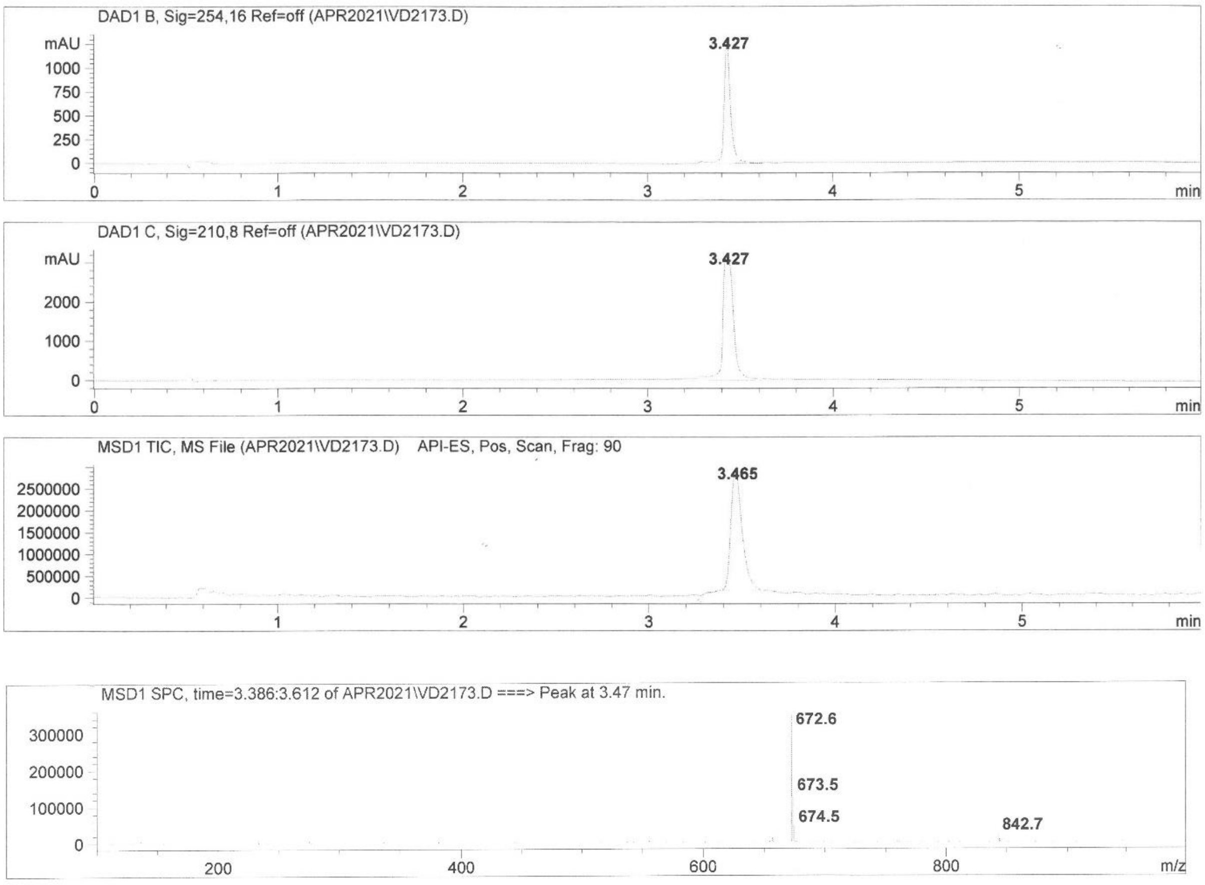

VD3152 (Compound 20)

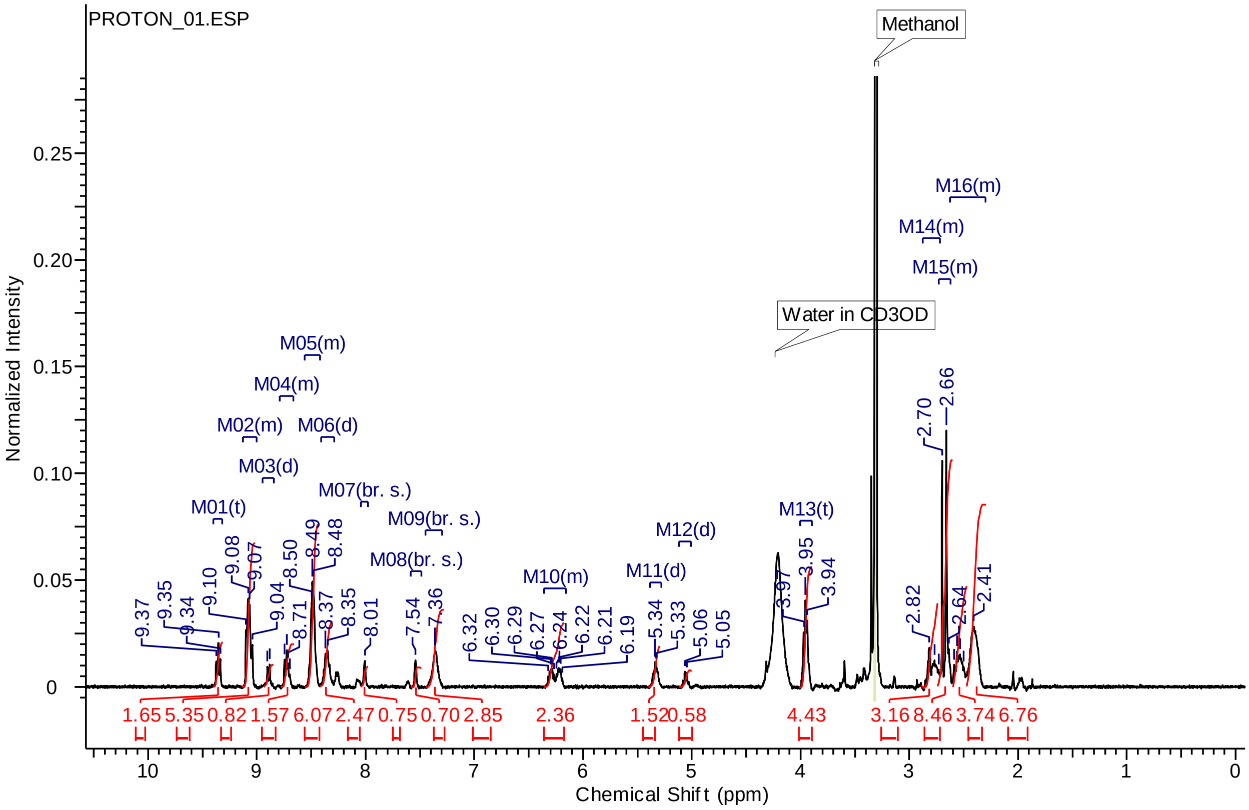


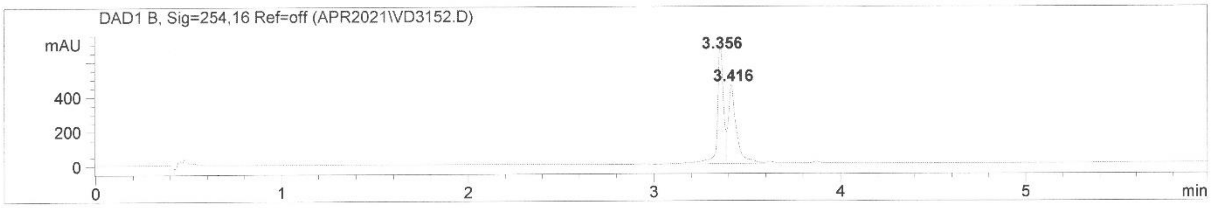


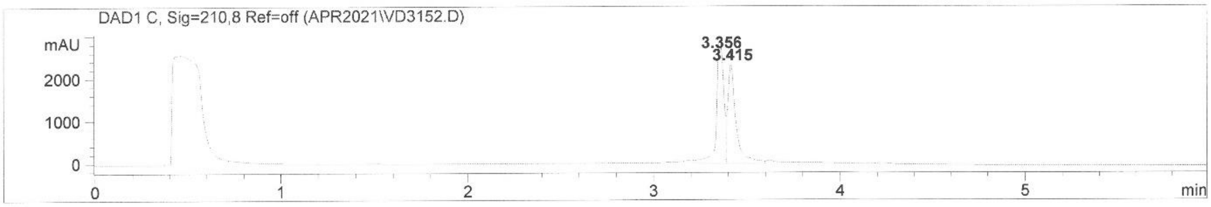


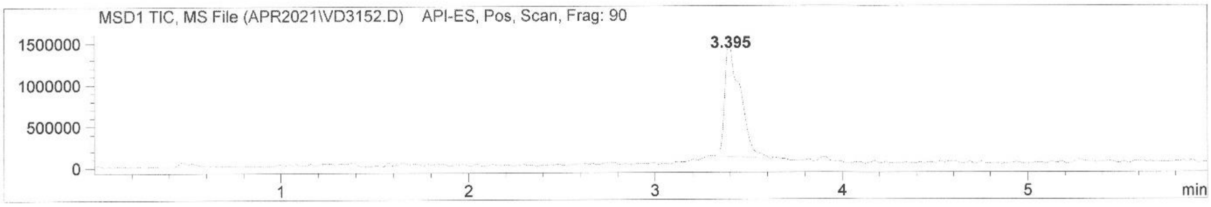


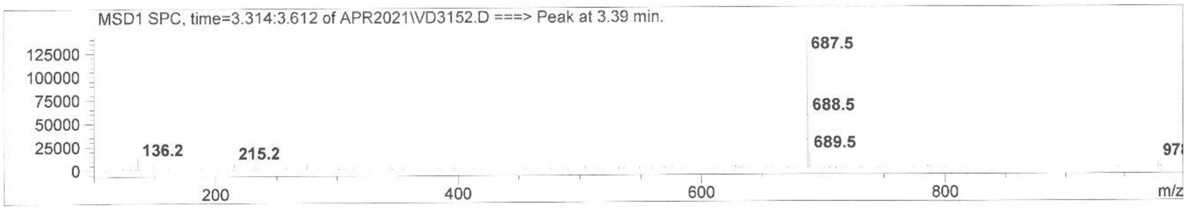

### VD3173 (Compound 19)

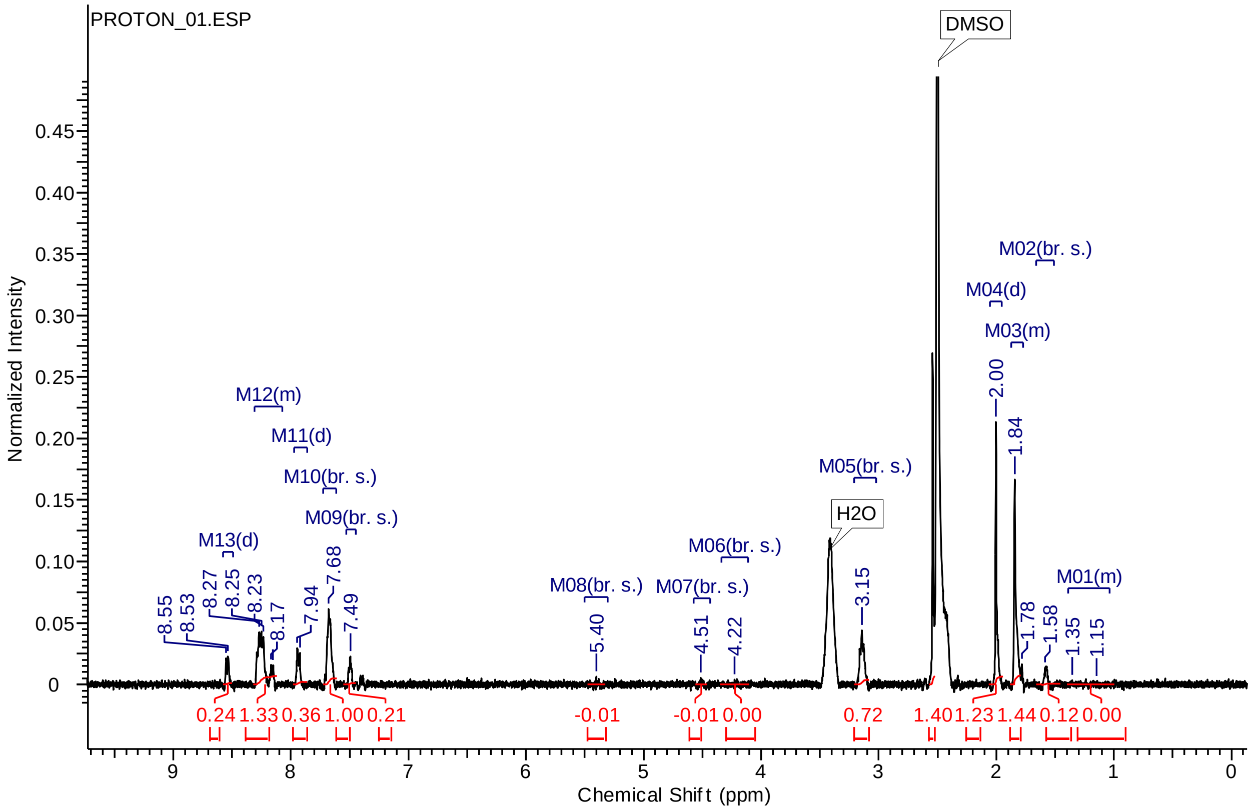


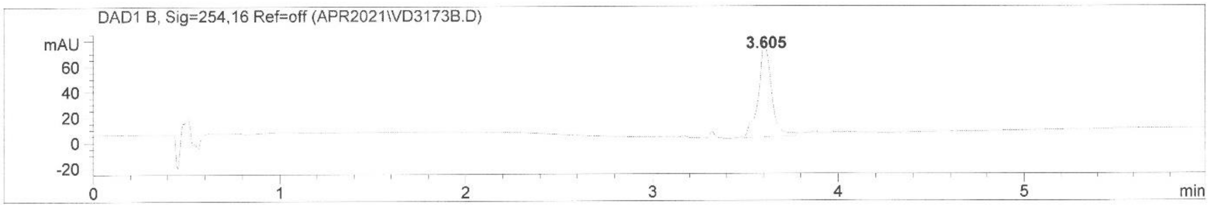


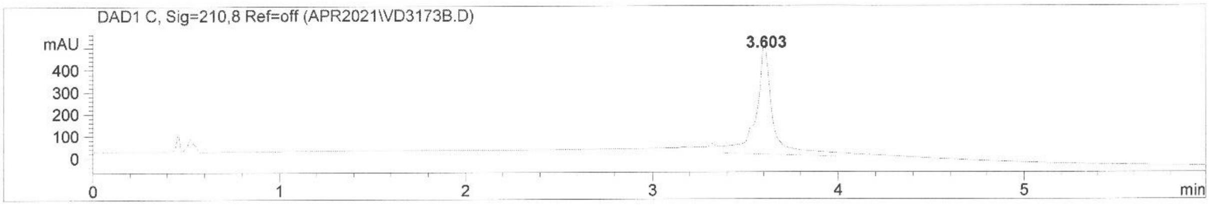


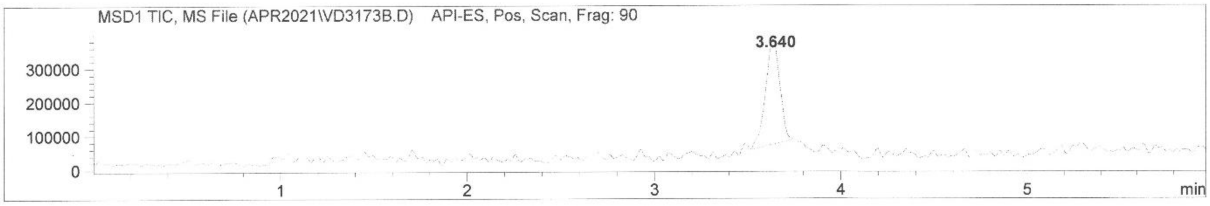


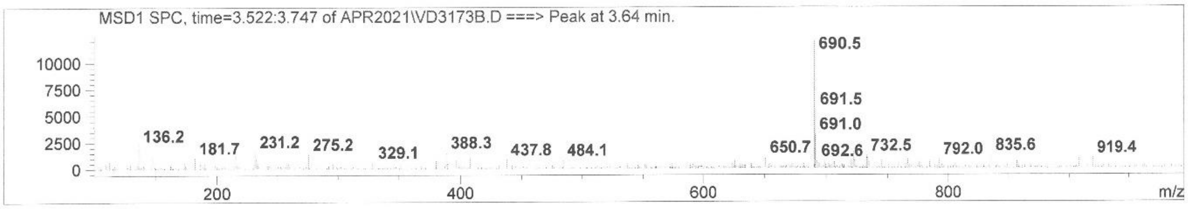

### VD4051 (Compound 21)

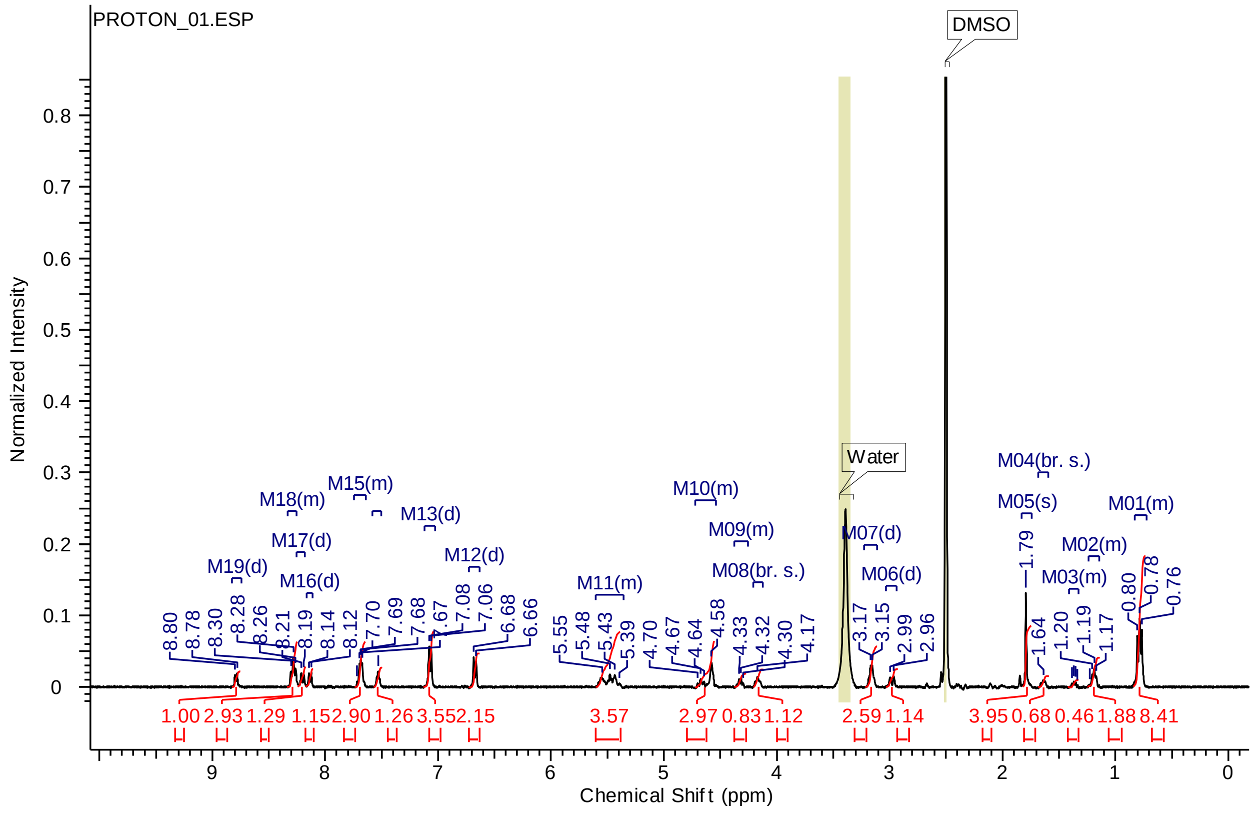


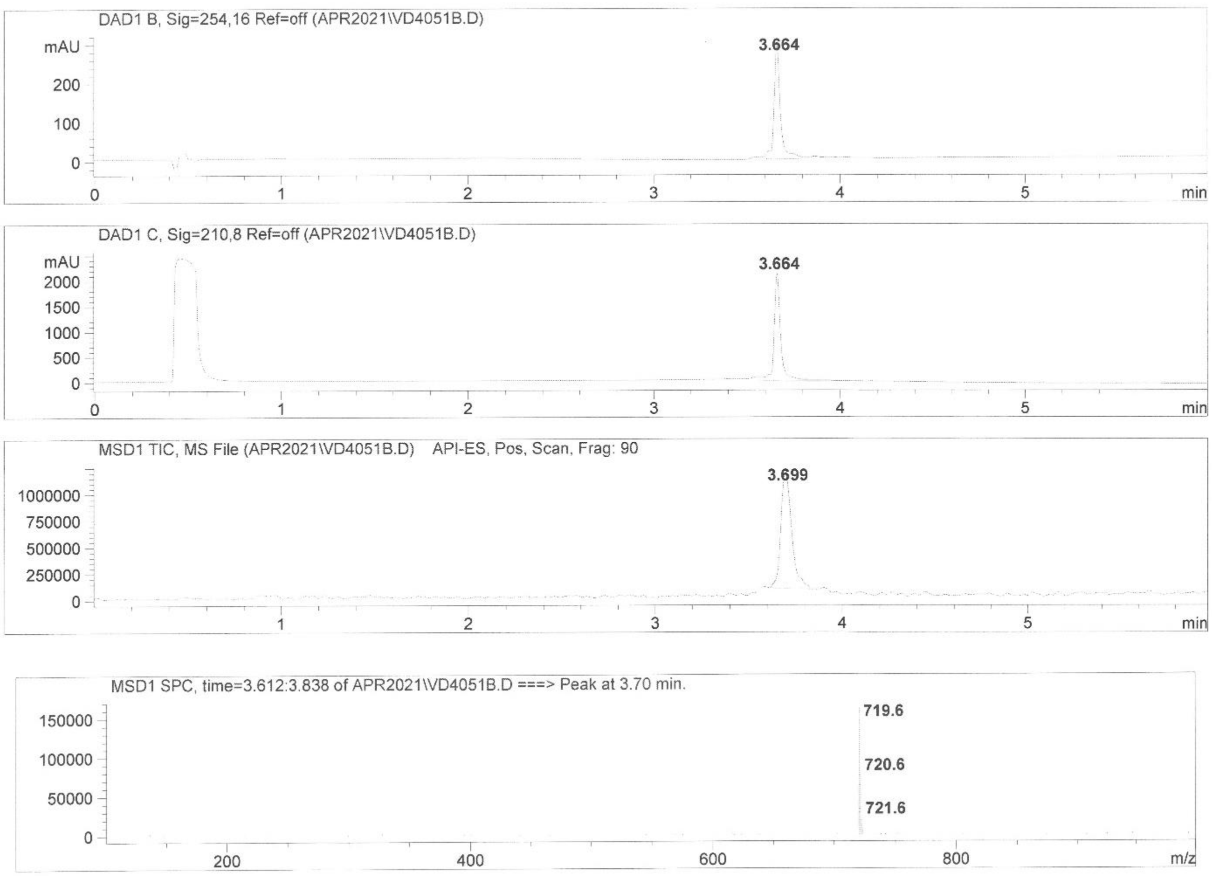

4. $K_m$ curve for Boc-QAR-AMC using full-length TMPRSS2 (Enzyme conc. 3 nM).

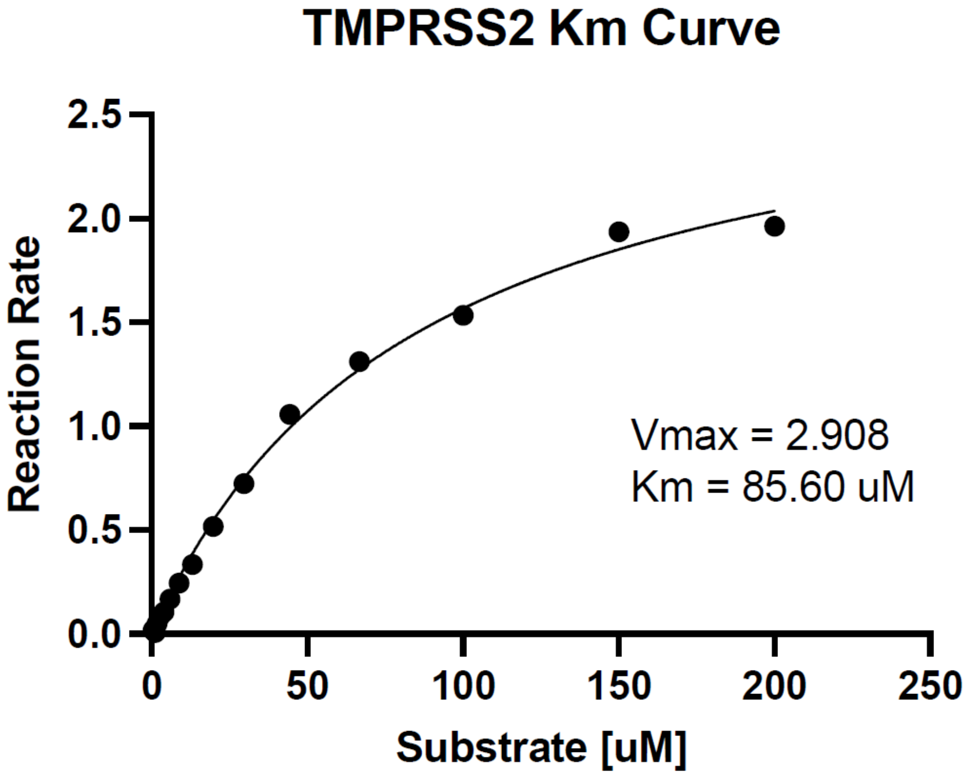

#### 5. IC$_{50}$ inhibition curves of full-length TMPRSS2/Boc-QAR-AMC (Table 1).

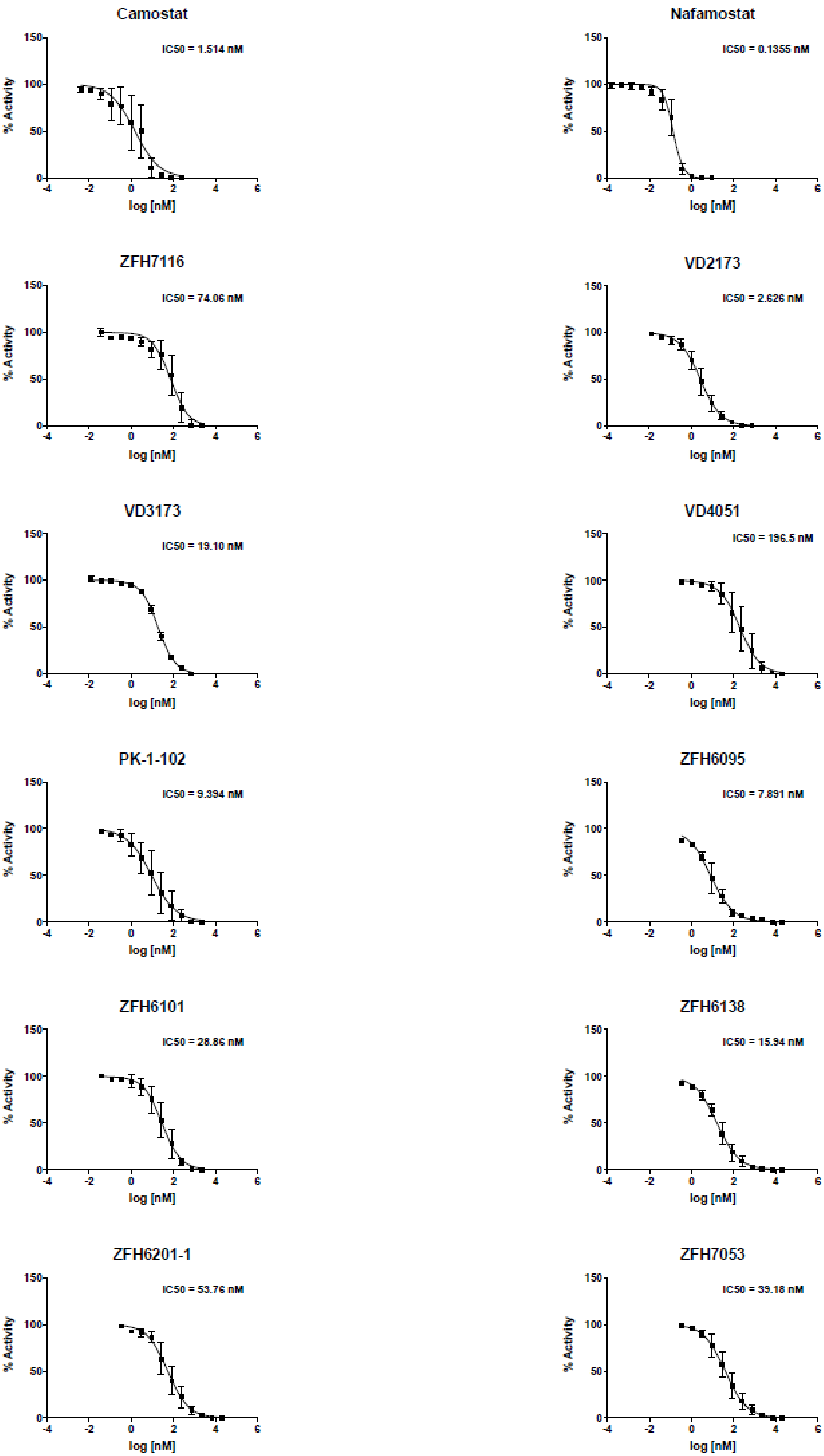

ZFH7063

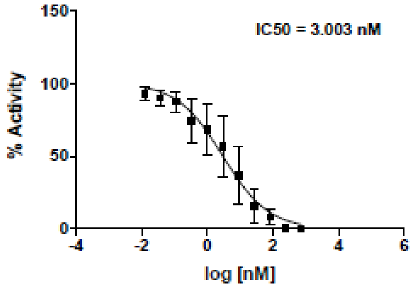


PK-1-89

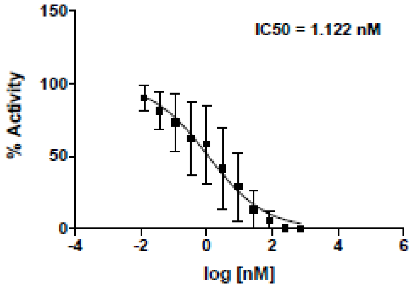


PK-1-18

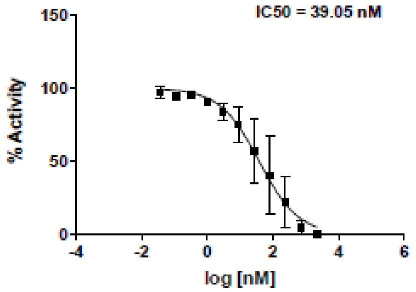


PK-1-45A1

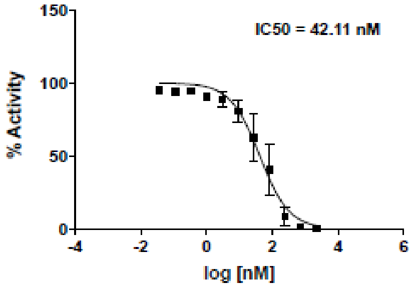


ZFH7064

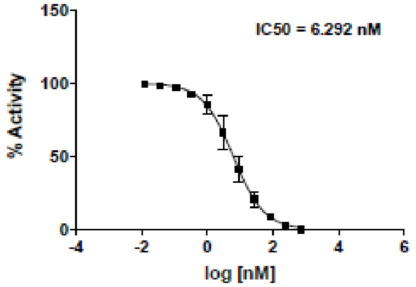


ZFH7006

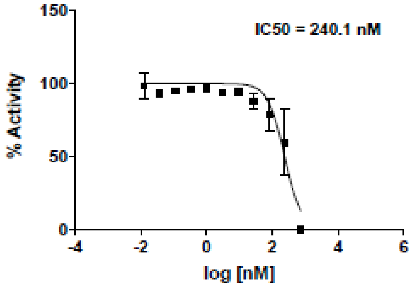


MM3116

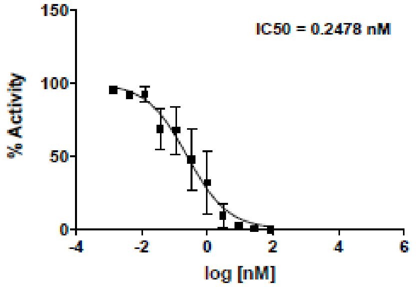


MM3144

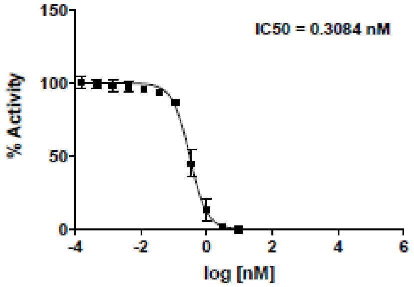


MM3123

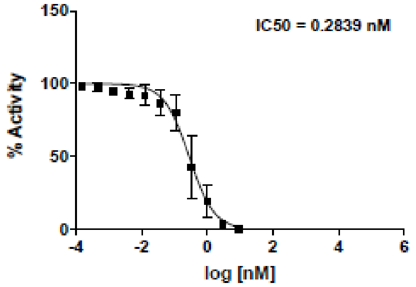


MM3122

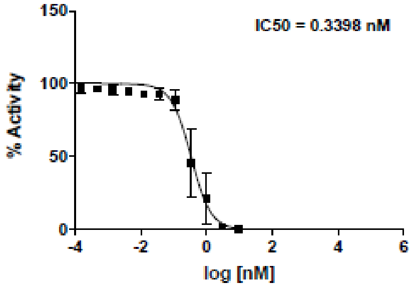

**6. Cell-based enzyme activity in HEK-293 cells. Acute toxicity of MM3122 (4) data. Activity of 1 and 2 and Camostat using Vero cells in pseudotype and chimeric VSV-SARS-CoV-2 viruses.**

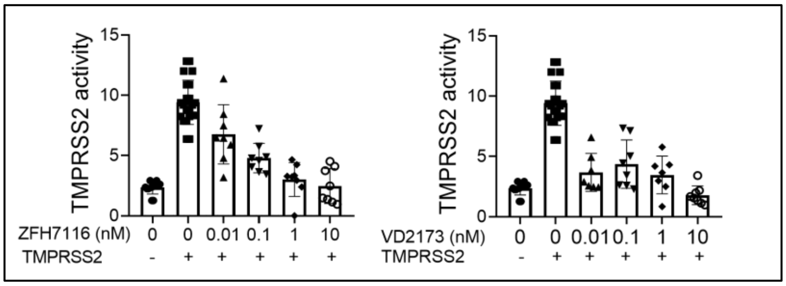


**Cell-based TMPRSS2 enzyme inhibition data of ZFH7116 (1) and VD2173 (2) in HEK-293 cells using Boc-QAR-AMC substrate.**

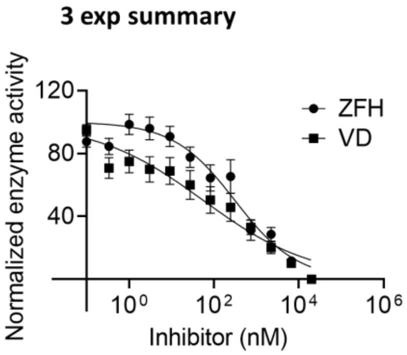


| | ZFH | VD |
|---|---|---|
| $IC_{50}$ (nM) | 314 | 57 |

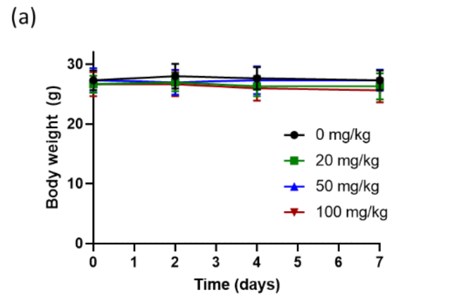


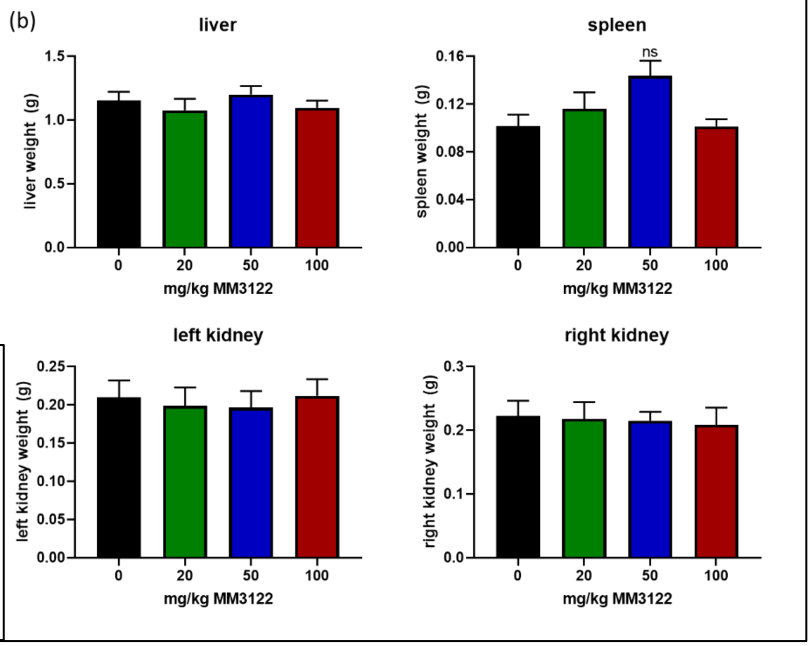


**MM3122 acute toxicity study.** NSG male and female mice (3 per group) were injected daily (IP) with 0, 20, 50, or 100mg/kg MM3122 for 7 days. No changes in (a) body weight or (b) liver, spleen, or left or right kidney weights were observed. Statistical analysis was performed using one-way analysis of variance (ANOVA) compared to control animals with Dunnett's multiple comparison test.

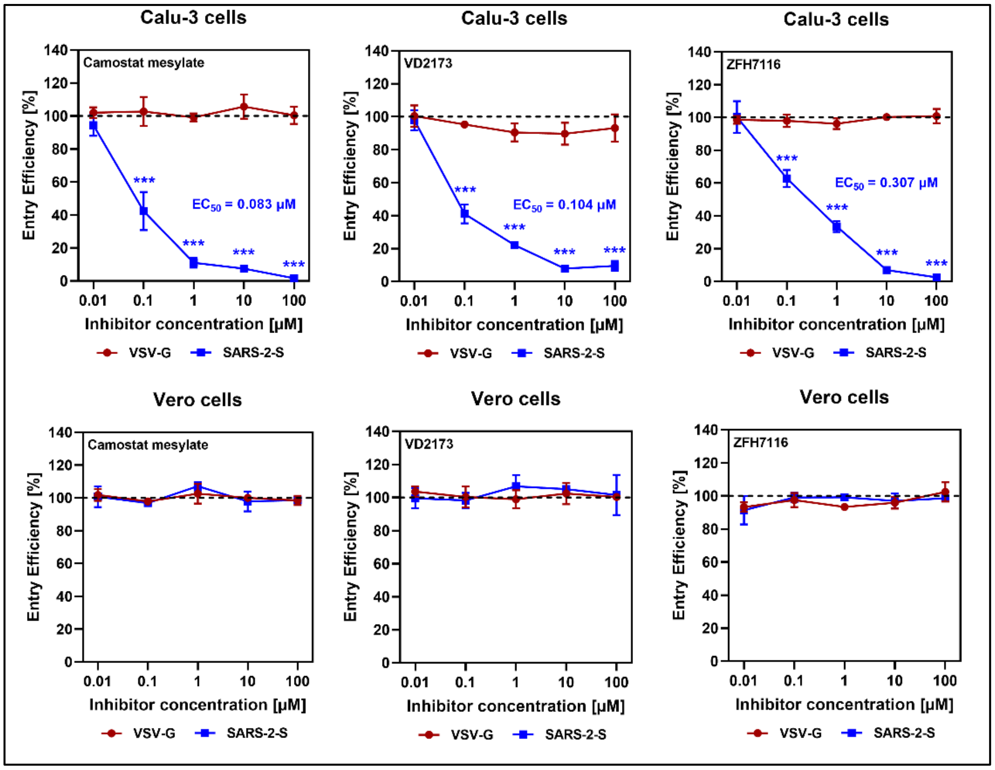


**Inhibition of SARS-CoV-2 cell entry into Calu-3 lung epithelial cells by ZFH7116 (1) and VD2173 (2) using VSV-SARS-CoV2-Spike protein Pseudotypes.** $EC_{50}$s are calculated from an average of 3 separate experiments. Camostat, an irreversible serine protease inhibitor, was used as a positive control.

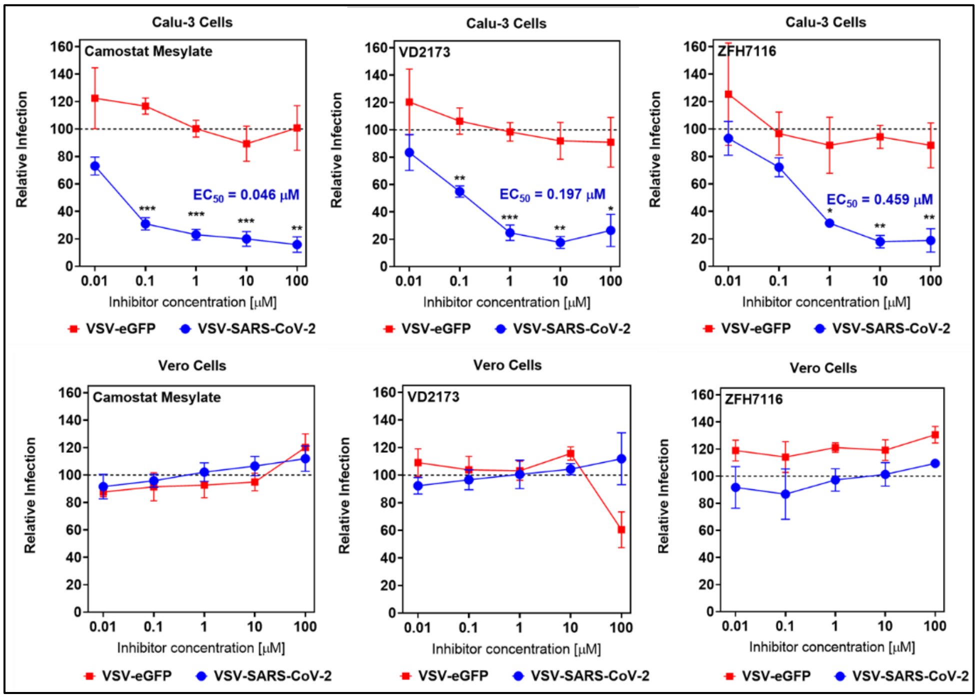


**Inhibition of SARS-CoV-2 cell entry into Calu-3 lung epithelial cells by ZFH7116 (1) and VD2173 (2) using VSV-SARS-CoV2-Spike protein Chimeras.** $EC_{50}$s are calculated from an average of 3 separate experiments. Camostat, a non-selective protease inhibitor, was used
